# Efficient 3D light-sheet imaging of very large-scale optically cleared human brain and prostate tissue samples

**DOI:** 10.1101/2022.07.14.500098

**Authors:** Anna Schueth, Sven Hildebrand, Iryna Samarska, Shubharthi Sengupta, Annemarie Kiessling, Andreas Herrler, Axel zur Hausen, Michael Capalbo, Alard Roebroeck

**Author notes:** contributed equally.

## Abstract

The ability to image human tissue samples in 3D, with both cellular resolution and a large field of view (FOVs), can improve fundamental and clinical investigations. Here, we demonstrate the feasibility of light-sheet imaging of ∼5 cm^3^ sized formalin fixed human brain and up to ∼7 cm^3^ sized formalin fixed paraffin embedded (FFPE) prostate cancer samples, processed with the FFPE-MASH protocol. We present a light-sheet microscopy prototype, the cleared-tissue dual view Selective Plane Illumination Microscope (ct-dSPIM), capable of fast, 3D high-resolution acquisitions, of cubic centimetre sized cleared tissue. We used Mosaic scans for fast 3D overview scans of entire tissue samples or higher resolution overviews of large ROIs with various speeds: a) Mosaic 16 (16.4 µm isotropic resolution, ∼ 1.7 hr/cm^3^), b) Mosaic 4 (4.1 µm isotropic resolution, ∼ 5 hr/cm^3^) and c) Mosaic 0.5 (0.5 µm near isotropic resolution, ∼15.8 hr/cm^3^). We could visualise ROIs around the border of human brain area V1/V2, and could demonstrate suitable imaging quality for Gleason score grading in prostate cancer samples. We show that ct-dSPIM imaging is an excellent technique to quantitatively assess entire MASH prepared large-scale human tissue samples in 3D, with considerable future clinical potential in prostate cancer.

## Introduction

Despite the clear advantages of large-scale microstructure visualisations, tissue samples in fundamental research and clinical pathology are still mostly examined with conventional light-microscopes in paper-thin tissue sections (ranging from approx. 5 – 100 µm), mounted on glass slides. This destroys the 3D organ structure and provides only limited 2D information over a small field of view (FoV). Therefore, significant advances are needed towards novel high-speed, high-volume, 3D multiscale microscopy approaches with sufficient resolution. This will allow for the detection of crucial details and overview features throughout the entire large tissue samples (ranging from mm to cm size).

The complex 3D structure of the human brain is inherently multiscale and exists of very small structures that extend over large distances, even entire brain areas [1]. The layered cortical cytoarchitecture, for instance, is defined by cellular density, size, and morphology at the microscopic scale, but its layers extend over entire cortical areas and hence layering occurs over centimetre scales. To enable quantitative characterisation, such as cell counting, throughout various layers and even entire brain areas, both large FoV overview scans need to be performed, as well as cellular high-resolution imaging. This kind of data is essential for e.g. realistic biologically informed neural modelling [2]. Therefore, the investigation of layered cytoarchitecture requires both high-resolution imaging and large FoVs.

In prostate cancer, tumours are characterised by multi-focality and a heterogeneous morphology with diverse histo-morphological patterns in 3D, throughout extended volumes. [3, 4]To date, a definitive diagnosis for prostate cancer requires histo-pathological verification of biopsies based on the Gleason Score grading [3]. This is challenging as shown by the inter-observer variability, which in turn can lead to under- or over-treatment of patients [4]. Furthermore, the criteria for “active surveillance” are determined by quantification of tumour extent and Gleason grade [5]. Since complete serial sectioning of prostate biopsy cores are rarely done, an under grading might occur in cases with multiple small foci of the prostate adenocarcinoma, because they are present at different levels in the paraffin blocks [5]. Moreover, false-negative biopsies may occur from the incomplete sectioning of tissue blocks. For instance, Paulk et al. demonstrate the occurrence of the prostate carcinoma in the deeper sections of the paraffin blocks that was absent on initial H&E sections. In current practice, routinely cutting through the entire paraffin blocks to increase tissue visualization may not be possible because of the higher workload and higher price, as compared with the standard procedure of sectioning only 3-4 levels.

In recent years, light-sheet fluorescence microscopy (LSFM), together with optical tissue clearing has been used for 3D visualisation and examination of rodent and human tissue at mesoscale to microscale resolution [6–11]. Various labs have developed optical clearing protocols, such as CLARITY [6, 8], iDISCO [9, 10] and CUBIC [12], which have mainly been applied to render mouse brains transparent, in order to understand both the brain’s structure and pathology [7]. However, the application of cleared tissue light-sheet imaging to large archival (i.e., fixed with aldehyde fixatives) adult *human* brain samples, in particular, has been a major challenge because of the sample size and the difficulties of applying clearing, labelling and imaging throughout large volumes of myelin-rich tissue [13].

Earlier work has demonstrated successful clearing, labelling and LSFM imaging of both human brain [7, 12–17] and human prostate cancer biopsies [18]. However, although human brain samples of between 1 mm^3^ [13] up to 1 cm^3^ [12] and 1.5 cm thick tissue slabs [14] were successfully cleared and labelled, the actual LSFM imaged sample size has been restricted to 1 mm^3^ [16] or 500 µm thick tissue slabs [13] with a reported maximum volume of ∼ 10.5 mm x 14.1 mm x 3 mm [17]. Zhao et al. [19] imaged the largest optically cleared and labelled human brain sample to date (7.5 x 5 x 0.4 cm) with confocal microscopy. However, a light-sheet microscopy set-up would increase the speed and scalability of the image acquisition by a large degree. Recent prostate studies, such as described and performed by Glaser et al. [18] focussed mainly on 1 – 2.5 mm core needle biopsies. An overview of the most prominent LSFM set-ups applied to human brain and prostate cancer samples (and other sample types) are listed in table 1, including sample volume and resolution. Both the Liu lab [18], as well as the Shroff lab [20], have mounted multi-immersion objectives for cleared tissue imaging to their open-top LSFM and the dual inverted Selective Plane Illumination Microscopy (diSPIM) respectively, to allow for cleared tissue imaging of e.g., human prostate and mouse brain samples. Although this shows that LSFM technology and clearing protocols are evolving rapidly, fast 3D cleared tissue imaging of large-scale human prostate and human brain samples of many centimetres in lateral size and many cm^3^ in volume has so far remained out of reach.

**Table 1:** Overview of various light-sheet fluorescence microscopy set-ups, including set-up name, use of human brain and/or sample type, sample volume and the imaging resolution. See reference for detailed information on author and research group.

| Reference | Name | Yes/No Human brain | Sample volume | Resolution |
| --- | --- | --- | --- | --- |
| [40] | ctASLM | No, mouse organs | Mouse brain, expanded mouse brain, average size of a mouse brain: $415 \pm 24$, 365 and 440 mm<sup>3</sup> | submicron, isotropic resolution, plus expansion, two resolution levels (~600 and ~300 nm, to 260 nm of axial resolution, submicron, isotropic resolution, high-speed aberration-free remote focusing, |
| [14] | Ultramicroscope and upright confocal microscope RSG4 from MAVIG | yes | 1.5 thick human brain slices (SHANEL) 2-4cm x 2-4 cm x 1.5.cm, Imaged by confocal: 7, 5 x 5 x 0.4 cm | Ultramicroscope: 4 μm lateral and 6.5 μm axial resolution. Small structures, such as axons (0.2 μm diameter) can be visualized.<br>Confocal: The RS-G4 is a true confocal system and thus provides the common advantages of such a system, like a high optical resolution and contrast (maximum resolution of 1024 x 1024 pixels at 5.9 frames per second).<br>(In general, a confocal microscope can reach below 1 μm resolution) |
| [12] | GEMINI system, custom-built light-sheet microscope combined with left and right light-sheet illumination units (developed by Olympus) and two macrozoom microscopes (MVX10, | yes | ~1 cm<sup>3</sup> tissue block of a post-mortem adult human cerebellum, imaged 1.5 mm thick | Ultramicroscope: 4 μm lateral and 6.5 μm axial resolution. Small structures, such as axons (0.2 μm diameter) can be visualized.<br>Custom-built LSFM: x-y: ~6.6 or 4.2 μm at |
| | Olympus) placed at the front and back sides of the sample. | | | $\lambda = 550$ nm, provided by Olympus, z: $\sim 10$ μm at $\lambda = 550$ nm, based on the effective N.A. for light-sheet illumination $\sim 0.03$ |
| [18] [41] | Open top light-sheet (OTLS), hybrid OPTLS with Non-orthogonal dual-objective (NODO) | No, prostatectomy slice, mouse organs, human breast cancer | $\sim 1$mm thick, core needle prostate biopsy, 2.5 mm thick prostatectomy slice, mouse organs, human breast cancer, 30 mm<sup>3</sup> tissue slices | sub-micron in-plane resolution at an imaging speed $\sim 1$ mm<sup>3</sup> min<sup>-1</sup> (per wavelength channel) with a maximum usable imaging depth of 0.5 cm and a lateral area of up to 10 × 10 cm, 1-mm diameter of all biopsies |
| [42] | diSPIM | No, C. elegans, etc | The C. elegans adult reaches a length of about 1 mm and a diameter of 80 μm. | isotropic spatial resolution (down to 330 nm) at high speed (200 images per s, 0.5 s for a dual-view 50-plane volume) |
| [15], [16], [13] | DIY confocal light-sheet (CLSM) and two-photon fluorescence microscopy (TPFM) and di2CLSM | Yes, mouse brain too | 1 mm<sup>3</sup> child, epilepsy); 500 μm thick slabs of human brain, whole mouse brain volume $\approx 223$ mm<sup>3</sup>, excised mouse cerebellum volume $\approx 73$ mm<sup>3</sup> | DIY confocal light-sheet (CLSM): the radial and axial point spread function (PSFs) are $(2.23 \pm 0.08)$ μm and $(9.1 \pm 0.4)$ μm respectively; di2CLSM: Our instrument, by fusing the two orthogonal views, enables an isotropic resolution of 1 μm along the three optical axes (x, y and z); TPFM: sub-cellular resolution |
| [11] | Ultramicroscope | Yes, human breast cancer sample | 30 × 20 × 5 mm (tumour tissue) | The resolution in XY ranges between 1-10 μm. Small structures, such as axons (0.2 μm diameter) can be visualized. |
| [17] | Light-sheet theta microscopy (LSTM) | Yes (may facilitate mapping of an entire post-mortem human brain (thick slab-by-slab) in a practical timeframe) | Human brain sample of ~ 10.5mm x 14.1mm x 3mm, expanded mouse brain 33.2 mm x 19.3 mm x 2 mm, rat brain ~ 2 cm wide and ~ 5 mm deep | LSTM achieves uniform axial resolution (~ 4–6 $\mu\text{m}$ FWHM) over the entire field of view |
| [7] | Ultramicroscope | Yes | 0.125 cm<sup>3</sup> AD human brain | 4 $\mu\text{m}$ lateral and 6.5 $\mu\text{m}$ axial resolution. Small structures, such as axons (0.2 $\mu\text{m}$ diameter) can be visualized |
| [43] | Scape | No, Zebrafish heart, C.elegans, mouse brain slice | The C. elegans adult reaches a length of about 1 mm and a diameter of 80 $\mu\text{m}$. | theoretical diffraction-limited lateral (x-y) and axial (z) resolutions of the SCAPE imaging geometry rival conventional light-sheet microscopy at 0.4-2 microns and 1-3 microns respectively over large fields of view for i.e., dendrites |
| [20] | diSPIM plus multi-immersion objectives, iSPIM, Zeiss Z.1, lattice light-sheet, confocal, widefield | No, mouse brain | average size of a mouse brain: 415 ± 24, 365 and 440 mm<sup>3</sup> | submicron resolution isotropic 0.79 ± 0.04 $\mu\text{m}$, |
| [44] | Tiling lattice light-sheets (LLS) in lattice light-sheet microscopy (LLSM), expansion | No, mouse brain), zebrafish embryo, c. elegans | ~3.8 x 3.8 x 7 mm<sup>3</sup> sample volume | Tiling light-sheet: Micron-scale (4 x 4 x 10 $\mu\text{m}^3$) to submicron-scale (0.3 x 0.3 x 1 $\mu\text{m}^3$) spatial resolution |

We demonstrate in this work that we optically clear, label, and image several cubic centimetres of human brain and prostate cancer (axial whole-mount sections) tissue samples. Previously, we presented MASH (Multiscale Architectonic Staining of Human cortex), a scalable clearing and labelling approach shown to be effective for up to 5 mm thick slabs of archival adult human brain tissue [21]. Here we processed Human brain samples were with MASH [21] clearing and labelling . Additionally, we introduce FFPE-MASH and for the first time successfully process FFPE human prostate samples. We then introduce the cleared-tissue dual view Selective Plane Illumination Microscope (ct-dSPIM) to perform efficient large cleared tissue LSFM 3D imaging, which far exceeds existing cleared LSFM human brain and prostate sample studies in imaged volume.

Our new approach allows for 3D imaging, visualisation and quantification of large volumes with high-speed, and an adjustable speed-resolution trade-off. To show the feasibility of this method, we describe LSFM imaging of the human brain and prostate. We use the ct-dSPIM to image the human brain (occipital lobe) up to ∼50 x 35 x 3 mm (>5 cm^3^) and human prostate cancer resections samples (the axial whole-mount section after prostatectomy) up to ∼40 x 35 x 5 mm (∼7 cm^3^). ct-dSPIM imaging allows for the extension of current methods and studies and enables the examination of several mm thick axial whole-mount prostate sections. The application of ct-dSPIM imaging to larger prostate cancer samples allows for novel 3D insights into both benign and neoplastic tissue morphology. This additional 3D knowledge on tumours can enhance tissue visualization throughout the block, lead to a better understanding of the prostate adenocarcinoma architecture and could possibly improve the diagnosis of prostate cancer.

## Materials and Methods

### Human brain samples

Human occipital lobe samples were taken from 3 human body donors (donor 1: male, 98 years; donor 2: female, 101 years; donor 3: female 90 years; no known neuropathological diseases, respectively) of the body donation program of the Department of Anatomy and Embryology, Maastricht University (UM). Tissue from donors 2 and 3 were used for the shrinkage evaluation (suppl. Fig. 2). The tissue donors had given their informed and written consent to the donation of their body for teaching and research purposes as regulated by the Dutch law for the use of human remains for scientific research and education (“Wet op de Lijkbezorging”). Accordingly, a handwritten and signed codicil from the donor posed when still alive and well, is kept at the Department of Anatomy and Embryology, UM, Maastricht, The Netherlands. The human brains were first fixed *in situ* by full body perfusion via the femoral artery. Under a pressure of 0.2 bar the body was perfused by 10 l fixation fluid (1.8 vol % formaldehyde, 20 % ethanol, 8.4 % glycerine in water) within 1.5-2 hours. Thereafter the body was preserved at least 4 weeks for post-fixation, submersed in the same fluid. Subsequently, brain samples were recovered by calvarian dissection and stored in 4 % paraformaldehyde in 0.1 M phosphate buffered saline (PBS) for 14-30 months.

### Human prostate cancer biopsies

All prostate resection specimen were retrieved from the archive of the Department of Pathology, Maastricht University Medical Centre (MUMC+) in the period between 2007 and 2015. For this publication three samples were used. This study was approved by the Medical Ethics Review Committee of the Maastricht University Medical Center in the Netherlands (2020-1537). All specimens were collected and studied in accordance with the protocol of the Dutch Code of Conduct for Observational Research with Personal Data and Tissue (2004) [22]. The specimens were received for diagnostic purposes and processed according to the internal standard operating procedures according to the national and international recommendations. The thickness of the axial whole-mount section of prostate specimen ranges between 3 and 5 mm. In short, the samples were initially fixated with 4% buffered formalin for 24 hours and further processed in the Vacuum Infiltrating Processor Tissue Tek VIP6 (Sakura Finetek USA, Inc, Torrance, CA, the USA), where the specimens were dehydrated by immersing in a series of ethanol solutions of increasing concentration until pure, water-free alcohol was reached. This step was followed by a clearing of the tissue in a Xylene solution, with consequent specimen infiltration paraffin. Finally, the specimens were embedded in paraffin according to the routine pathology diagnostic procedures in the HistoCore Arcadia Embedding Center (Leica Microsystems B.V., Amsterdam, the Netherlands).

### Optical clearing and labelling with MASH and FFPE-MASH

All samples were cleared and labelled with neutral red using the MASH-NR protocol as described in our previous publication [23]. Since the standard prostate clinical protocol produced formalin fixed paraffin embedded (FFPE) samples, a modified MASH protocol was used (FFPE-MASH-NR) to process FFPE prostate samples. In the FFPE-MASH-NR protocol prostate samples had to be deparaffinised before standard MASH clearing. For this purpose, samples were incubated for 3 – 7 days in Xylene depending on sample size (for each sample a Xylene volume of about 200 ml was used). Paraffin blocks were manually trimmed as much as possible before the incubation. After that, samples were rehydrated 1h each in 100 ml Xylene, 2x 100 %, 70 %, 50 % ethanol (EtOH) and finally PBS. Rehydrated samples were kept in 4 % buffered PFA solution until use. Clearing and labelling of the samples with FFPE-MASH followed the same steps described below for the brain samples. During all these steps, prostate samples were kept in individual glass containers and incubated in at least 50 ml of the respective solution. Containers were kept in a shaker at all times.

All samples were cleared and labelled with neutral red using the MASH-NR protocol as described in our previous publication [23]. In short, samples were dehydrated for 1h each in an aqueous mixture (v/v) of 20 %, 40 %, 60 %, 80 %, and 100 % methanol (MeOH) at room temperature (RT) and 1h in 100 % MeOH at 4 °C. After that, samples were bleached overnight in a freshly prepared, chilled solution of 5 % H2O2 in MeOH at 4 °C. Samples were then rehydrated for 1 h each in 80 %, 60 %, 40 %, 20 % MeOH and permeabilised in phosphate buffered saline + 0.2 % (v/v) Triton X-100 pH 7.4. This was followed by an 1h incubation in freshly filtered aqueous solution of 50% potassium disulfite (w/v) and 5 quick rinses followed by 1h washing in distilled water. Labelling for cytoarchitecture was performed for 5 days in a solution of 0.001 % neutral red in phosphate-citrate buffer (aka McIlvain buffer)[24] at pH 4. Samples were flipped after half the incubation time. After labelling, samples were washed 2x1h in McIlvain buffer pH 4 and dehydrated for 1h each in 20 %, 40 %, 60 %, 80 %, 2x 100 % MeOH. Delipidation was performed overnight in 66 % dichloromethane (DCM)/33 % MeOH, followed by 2x 1 h 100 % DCM. Finally, samples were incubated with ethyl cinnamate (ECi) as refractive index matching solution (RIMS). All steps were carried out at RT.

For the incubation of multiple coronal slices of whole occipital lobes, a glass jar with a diameter of 8 cm was used, with spacers made from polyethylene or polytetrafluorethylene, in order to provide compatibility with the organic solvents used during delipidation. To prevent the plastic spacers from leaving impressions on the tissue, pieces of filter paper were placed above and below the tissue. The glass jar was filled completely for each step described above (a volume of approx. 200 ml) and the solutions were constantly agitated with a magnetic stirrer during the entire processing.

### Evaluation of tissue shrinkage

In order to evaluate the tissue shrinkage induced by MASH, we measured the surface area of a total of 13 brain slices from 3 different donors (sample “Occlobe16” n=4, tissue from the same donor as shown in figures 3-7; sample “Occlob10” n=6; sample “Hemi6” n=3, coronal slices anterior to the occipital lobe). “Occlobe10” slices (n=6) were taken from another lobe that has been imaged for an independent project. “Hemi6” slices (n=3) were taken from pre-sectioned material that was provided by the Department of Anatomy and Embryology at Maastricht University. These manually cut coronal brain sections of approx. 1 cm thickness were further sliced into 5 mm thick samples, before MASH processing. Scaled digital images were taken before clearing and at the end of the clearing procedure, after RI-matching. The circumference of the tissue slices was manually segmented in FIJI and the area before and after clearing measured (suppl. Fig. 2).

### Imaging chambers

For ct-dSPIM imaging, large customized (20 x 15 x 3 cm, volume of approx. 900 mL) imaging chambers (Fig. 1) were 3D printed in either ECi resistant watershed material (Somos®WaterShed XC 11122) or polypropylene (PP). The printing was performed either by SKM Rapid Modelling b.v. (Helmond, the Netherlands) via stereolithography (SLA) for the watershed prints or produced via Selective Laser Sintering (SLS) by Materialise NV (Leuven, Belgium) for the PP chambers. The imaging area of the chambers was equipped with a 178 x 127 x 1.2 mm glass slide (Ted Pella Inc., Redding, US) as a bottom to decrease light reflections and glued with pure silicone sealant. All samples were glued onto the glass slide in the 3D printed imaging chamber with hot glue (Rapid AB, Hestra, Sweden). Then, the imaging chamber was filled with at least 500 ml ECi solution (RI 1.56) with larger quantities depending on sample size.

**Figure 1:**
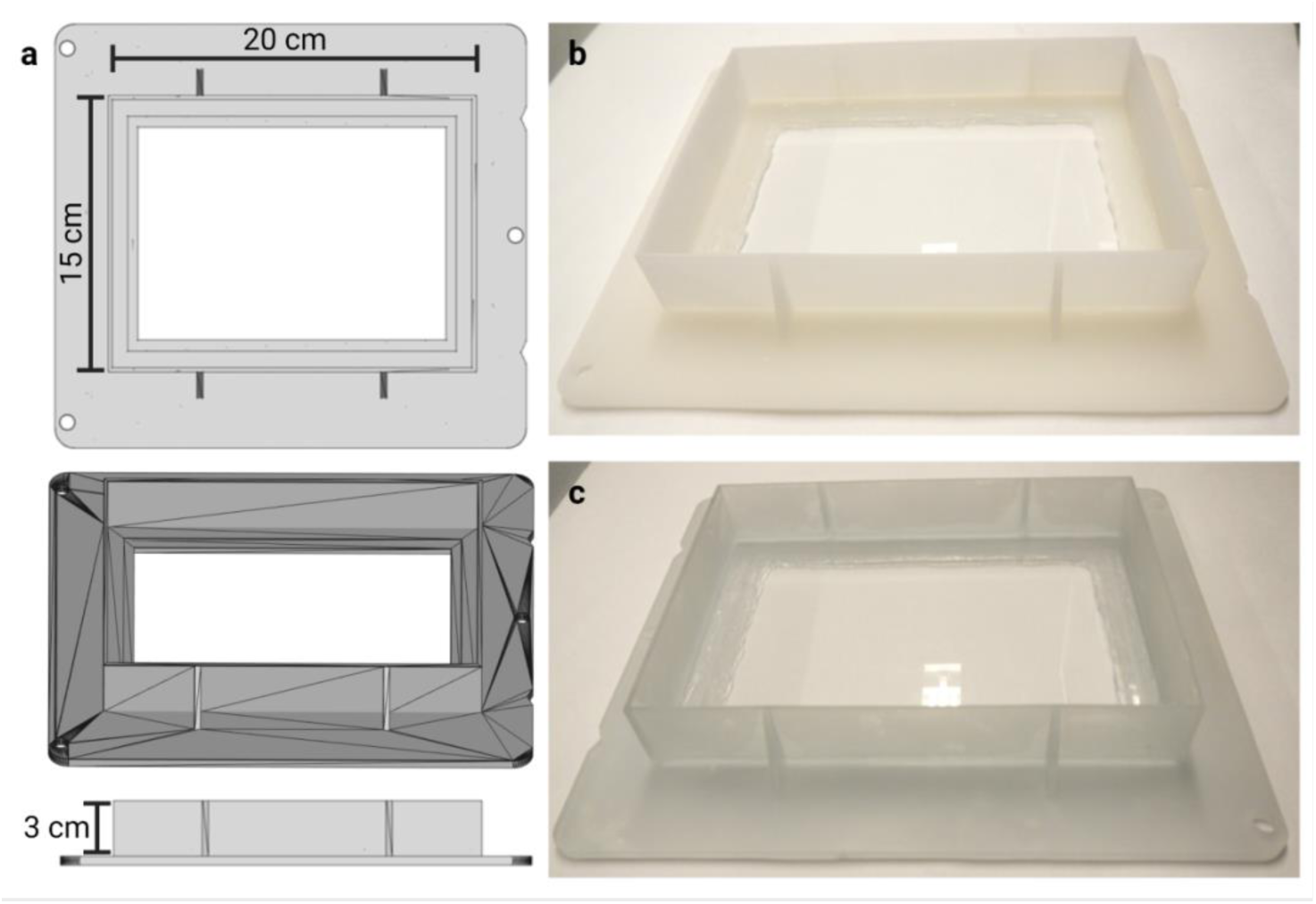
3D printed imaging chambers for large, cleared samples. **a**) 3D rendering of the larger imaging chamber used routinely on the ct-dSPIM set-up. The chambers have a volume on 0.9 l (20 cm x 15 cm x 3 cm). Chamber is shown in top view (top), oblique (middle), and side view (bottom). **b**) Chamber prototype printed with SLS in PP. **c)** Chamber prototype printed with SLA in Somos®WaterShed XC 11122. Samples are mounted on top of glass slides, which are sealed with silicon to prevent the chamber from leaking RIMS.

### ct-dSPIM light-sheet microscope set-up for large-scale cleared tissue imaging

The ct-dSPIM microscope is aimed at 3D imaging of very large, cleared tissue samples with a thickness of up to 5mm and a lateral size limited only by XY-stage travel and imaging time limits. It was derived from the diSPIM (dual view inverted Selective Plane Illumination Microscopy) system [25] and co-developed together with Applied Scientific Instrumentation Inc. (ASI, Eugene, US). An optical schematic of the ct-dSPIM system is shown in Figure 2. The laser light-source (Coherent obis, laser line 552nm LS 40mW LASER SYSTEM: FIBER PIGTAIL: FC) has a single-mode fibre output with a numerical aperture (NA) of 0.12. The emitted laser light is collimated and passes through an electronically tuneable (ETL) lens (C 60-TUNELENS-100, ASI) into the light-sheet “scanner” (MM-SCAN-1.2, ASI). The tuneable lens allows the axial position of the beam waist at the sample to be electronically controlled. The scanner contains a 2D micro-electro-mechanical mirror (MEMS) which sweeps the Gaussian beam across the sample once per camera image to form a “virtual” or “digital” light-sheet. The other axis of the MEMS mirror is used to adjust the l coincident with the detection objective’s focal plane. The scanning beam is relayed to the back focal plane of the multi-immersion illumination objective (#54-10-12, Special Optics/ Applied Scientific Instrumentation (ASI).

**Figure 2:**
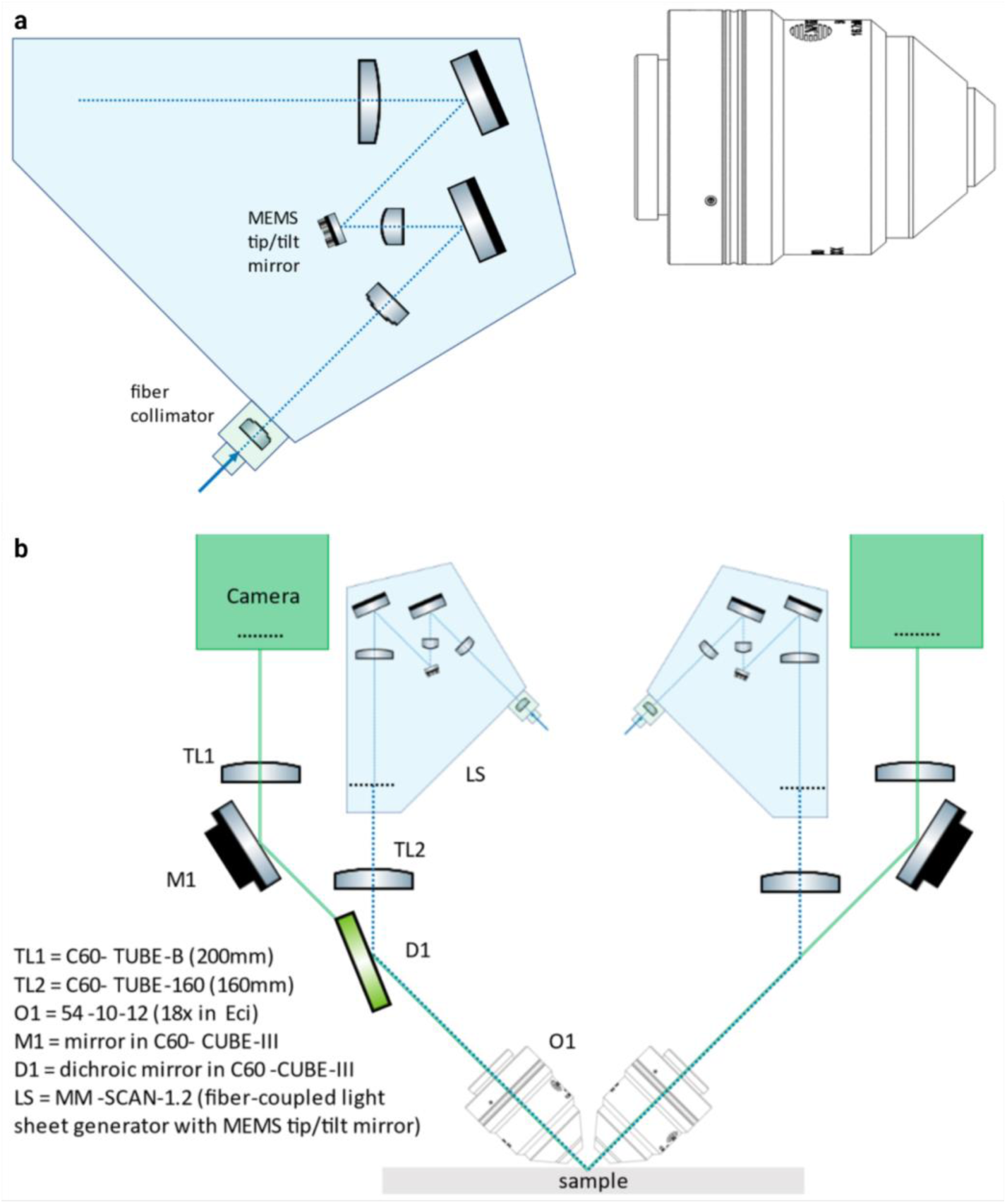
Optical layout of the ct-dSPIM. **a**) The emitted laser light is collimated and passes through an electronically tunable (ETL) lens (C60-TUNELENS-100, ASI) into the light-sheet “scanner” (MM-SCAN-1.2, ASI). The ETLf allows the axial position of the beam waist at the sample to be electronically controlled. Legend continued on the next page. The scanner contains a 2D MEMS mirror, which sweeps the Gaussian beam across the sample once per camera image to form a “virtual” or “digital” light-sheet. The other axis of the MEMS mirror is used to adjust the light-sheet coincident with the detection objective’s focal plane. **b**) The imaging paths are the two possible light paths (path A and B). For each path, the scanner is on one side, and the imaging piezo and camera are on the opposite side. The light travels through the different components, such as the dichroic mirror and emission filter, as depicted (excitation paths: blue dotted line; emission paths: solid green line). The filtered fluorescence is focused onto a 2048 × 2048 pixel sCMOS camera (ORCA-Flash4.0 V3, Hamamatsu) by a tube lens (C60-TUBE-B, ASI; f=200 mm).

Fluorescence is collected by an identical multi-immersion detection objective (#54-10-12, Special Optics/ ASI) and results in ∼1 μm resolution laterally (depending on the RI of the imaging solution). The objectives are compatible with a refractive index (RI) range from 1.33 to 1.56 and a working distance (WD) of 12 mm. Both the numerical aperture (NA) and effective focal length (EFL) vary with refractive index, but for ECi the NA is ∼0.43, the EFL is 11.2 mm, and the magnification is 17.9x. The maximum imaging depth is 5 mm and is limited by the physical clearance of the two objectives. The objectives are held by mechanics that include manual fine adjustments for co-aligning the two objectives (SPIM- DUAL-K2, ASI).

The excitation and detection paths are combined on a polychroic mirror (ZT488/543/635rpc-UF2, Chroma) and a motorized filter wheel (FW) (FW-1000-8, ASI) with three emission filters (ET519/26m; ET576/31m; ET655lp, Chroma), which allows three-color imaging. In this work, the aim was efficient single-color imaging over very large volumes and only the middle color band with ET576/31m emission filter was used. The filtered fluorescence is focused onto a 2048 × 2048 pixel scientific Complementary Metal–Oxide–Semiconductor (sCMOS) camera (ORCA-Flash4.0 V3, Hamamatsu) by a tube lens (C60- TUBE-B, ASI; f=200 mm). The tube lens provides a Nyquist sampling of 0.3625 μm/pixel at an RI of 1.56, with a horizontal field of view of 0.74 mm over the 2048 pixels of the camera.

Image strips are collected with a combination of stage-scanning (i.e., the sample is moved through a stationary lightsheet using the XY stage, Suppl. Fig. 1), lateral/vertical tiling using a motorized XY stage (MS-8000, scan-optimized) and motorized Z actuators (Focusing Translation Platform (FTP-2050), ASI). The acquisition speed is >10^8^ voxels/sec at an exposure time of 10 ms. The stage-scanning firmware emits an internal (time-to-live) TTL trigger which ensures the reproducible start positioning (<1 μm) of each image strip. An ASI tiger controller (TG-1000) controller contains control electronics for the motorized stages, MEMS mirror, tunable lens, and camera and laser triggers. It synchronizes all these elements with sub-ms precision during each image strip based on the initial stage-scanning trigger.

The microscope is controlled by µManager 1.4.22, a free open-source microscope control software[26]. The ASI diSPIM plug-in in µManager is used to both align the microscope and to setup and perform acquisitions. The stage control plug-in is used to make static ETL adjustments on both ETL’s (V, left; W, right).

For some applications, the 3D information from a single view or stack is sufficient. However, for the large sample imaging targeted here, extensive tiling is used along y-z Mosaic acquisitions, usually in combination with long image stacks stage scanned in the x direction (Suppl. Fig. 1). As a dual view system, the ct-dSPIM has the further advantage that the role of the two objectives can be reversed to collect another stack from a perpendicular direction at the expense of twice the imaging time. However, since the emphasis here is on > 1 um resolution fast large volume imaging, the dual view imaging mode is not employed in this work.

### Image acquisition and multiscale scans

The MASH-NR labelled human brain and prostate tissue sample were imaged with the 552 nm laser line at 1mW (OBIS) and with an exposure time of 10 ms throughout. For both samples, an overview scan of the entire cubic centimetre-scale tissue slices was performed with 16.4 µm isotropic sampled resolution (which we name a Mosaic 16 acquisition). Additionally, for the human brain sample, we have performed a multiscale scan consisting of: a) a Mosaic 16 overview scan of the whole tissue slice, b) an acquisition for a large selected FoV (∼ 15 x 17 x 3 mm) with 4.1 µm isotropic sampled resolution (termed Mosaic 4) and c) an acquisition for a smaller FoV (∼5 x 10 x 3 mm) with high-resolution (0.725 µm x 0.5127 µm x 0.5127 µm) for specific region of interests (termed Mosaic 0.5). The imaging speed is different depending on which Mosaic imaging is deployed. For fast Mosaic 16 overview scans the imaging speed is at 1.7 h /1cm3 and for higher resolution ROI Mosaic 4 and Mosaic 0.5 scans the imaging speed is at 5h/ 1cm3 and 15.8h/1cm3, respectively (Table 2).

**Table 2.** Resolution, imaging speed and slice step for Mosaic 16, 4 and 0.5 ct-dSPIM scans, respectively.

|  | Isotropic Resolution | Speed | Slice step |
| --- | --- | --- | --- |
| Mosaic 16 | 16.4 $\mu\text{m}$ | 1.7hrs/1cm<sup>3</sup> | 11.60 $\mu\text{m}$ |
| Mosaic 4 | 4.1 $\mu\text{m}$ | 5 hrs/1cm<sup>3</sup> | 2.90 $\mu\text{m}$ |
| Mosaic 0.5* | 0.5$\mu\text{m}$<br>* (0.725 $\mu\text{m}$ x 0.5127 $\mu\text{m}$<br>x 0.5127 $\mu\text{m}$) | 15.8 hrs/1cm<sup>3</sup> | 0.363 $\mu\text{m}$ |

Whereas the Mosaic 16 scans provide 3D datasets of an entire tissue block at 16.4 µm isotropic mesoscopic sampled resolution, the Mosaic 4 and Mosaic 0.5 show regions of interest at 4.1 µm and nearly 1 µm isotropic sampled resolution, respectively. The slice steps at acquisition are 11.60 µm for Mosaic 16, 2.90 µm for Mosaic 4, which lead to isotropic sampled resolutions after deskewing due to √2 scaling (Suppl. Fig. 1). The raw datasets were further downsampled as follows: For Mosaic 16, 16x in plane (32x45 pixels), and for Mosaic 4, 4x (128x186 pixels), to match the step size of the microscope and produce an isotropic dataset of 16.4 µm (Mosaic 16) or 4.1 µm (Mosaic 4) isotropic after deskewing. For the Mosaic 0.5, the slice step was 0.363 µm and in-plane downsampling of 2x2 was performed. With these parameters the Mosaic 16 scans are acquired at the highest possible speed, limited only by maximum stage scanning-speed. The Mosaic 4 scans are acquired close to a Nyquist sampling of the ∼8 µm axial optical resolution (i.e., theoretical light sheet thickness), and the Mosiac 0.5 scans are acquired close to a Nyquist sampling of the ∼1 µm lateral in-plane optical resolution. When acquiring image volumes with the ct-dSPIM, which uses stage scanning to move the sample through the imaging plane (i.e. the light sheet), the axes do not correspond to orthogonal XYZ coordinates. Instead, the camera Z-axis is at a 45° angle towards the stage scanning direction (Suppl. Fig. 1). Therefore, stage-scanned image acquisition leads to a skewed parallelepipedal stack shape with non-orthogonal axes, which will be warped when viewed in the orthogonal axis system. A deskewing operation transforms the skewed parallelepipedal stack into a rectangular stack with the traditional orthogonal axes (Suppl. Fig. 1b). The axis labeling used in figures in this work is the image viewer axis labeling with the x axis along the 3-5mm thickness of the tissue and the z axis along the scanning direction of the image stacks (Suppl. Fig. 1c).

### Data processing and visualisation

Down-sampling is performed by scaling the images to the final pixel size with the scaling function in Fiji [27]. The down-sampling reduced the size of the data set to ∼3-4 GB for a Mosaic 16 scan of an entire occipital lobe slice. This down-sampled data was further processed with the FIJI PlugIn BigSticher [28]. First, the data was deskewed using the “(de)skewing” option and then stitched via the Phase Correlation option, without further down-sampling. The stitched data was then resaved as a 16-bit tif file. In some cases, for visualization, the final isotropic, fused data set was resliced in FIJI along the YZ direction to provide a coronal view on the samples. The overall down-sampling time for both a Mosaic 16 and Mosaic 4 scan takes 12h on a PC workstation with 32 RAM, Intel(R) Xeon(R) CPU E5-1650 v3 @ 3.50GHz and 10 HDD storage. Moreover, for a Mosaic 0.5 no down-sampling is needed. The total stitching time with the BigStitcher PlugIn takes approximately 1h for the Mosaic 16 and 2h for the Mosaic 4 scan, respectively. For the Mosaic 0.5 approximately 3 days of stitching time are needed in BigStitcher.

3D visualisation and cell count of volume stacks were performed with arivis Vision4D software (version 3.4.0). For this purpose, a single stack from a Mosaic 0.5 acquisition in human brain area V2 and covering all cortical layers, was used. The deskewed, stitched, and resliced stack was processed with an automated pipeline in Vision4D. The raw data was first filtered with the morphology filter option and cells were segmented with the “blob finder” segmentation method. To derive cell numbers per layer, the cortical layers and white matter were manually segmented with the polygon selection tool over the entire volume. Layer III was divided in two sublayers IIIa and IIIb based on notable density and cell size differences (e.g., large pyramidal neurons appearing in layer IIIb). A second pipeline was then applied to create compartments based on the manually segmented layers, which included only the segmented objects from the cell segmentation pipeline that were fully contained within the layer segment borders and had a size of larger than 125 µm^3^. Features derived from these compartmentalised segments were then extracted into itemized .csv tables.

To visualise the isotropic resolution of the lower resolution Mosaic 16 and Mosaic 4 datasets, orthogonal views were created in FIJI [27] by reslicing the data and creating MIPs for each axis. Videos were created both with FIJI as well as with Vision4D. Figures were created with biorender (https://www.biorender.com).

### Manual cell count validation

In order to validate the automated cell segmentation pipeline, manual cell counting was performed on the same dataset. The data volume was divided into a 100x100 µm grid, extending over the entire depth of the stack, in the Vision4D environment with a custom-made python script. Subsequently, 10 planes and 5 different ROIs per plane were pseudo-randomly selected using the “randi” function in MATLAB (R2015b, MathWorks Inc.) as a random number generator. Cells were manually counted within each ROI, considering only cells that were either fully contained within the ROI boundaries or intersecting with the left and/or lower border of the 100x100 µm box (supplementary Figure 3A). Violin plots of the data distribution were created in MATLAB using the script by Bechtold (2016; DOI: 10.5281/zenodo.4559847) for the manually counted cells, the unfiltered “blob finder” segmentation results, as well as the filtered results containing only segments larger than 125 µm^3^ (“cell bodies filtered”; supplementary Figure 3B).

## Results

### Large-scale imaging of optically cleared human tissue samples

We have performed multiscale 3D imaging on MASH prepared human brain (Fig. 3-7) and the axial whole-mount prostate sections (Fig. 8, 9, suppl. Fig. 3, 4) with a large FoV with Mosaic 4 and Mosaic 16 acquisitions and a moderate FoV and high-resolution for a specific region of interests. Mosaic 16 of human brain samples (Fig. 3, 4, 6) allowed for relatively high-speed (approx. 8-16 hrs, ∼1.7 cm^3^/h, 16.4 µm isotropic, Table 2) overview scans of an entire tissue block up to 5 cm x 3 cm in lateral size and 3 mm thick. Smaller ROIs of the human brain samples were imaged with a Mosaic 4 (Fig. 5, 6). and Mosaic 0.5. (Fig. 7). We present and discuss these results in turn.

**Figure 3:**
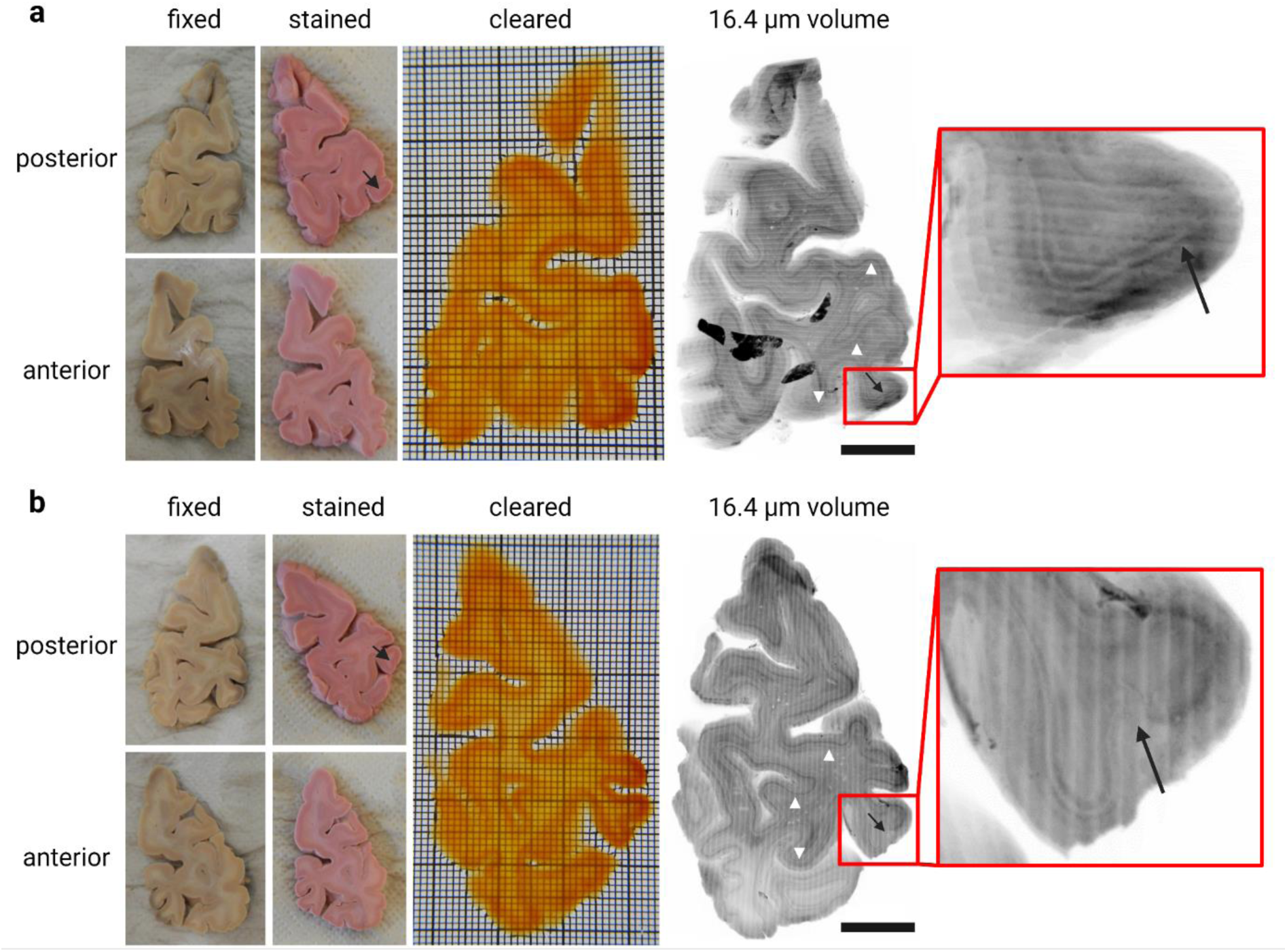
MASH processing and Mosaic 16 acquisition of 3 mm thick human occipital lobe sections. **a**) Sample taken approx. 6 mm anterior to occipital pole (third 3 mm thick section in posterior to anterior direction). From left to right: Formalin fixed, unstained sample and MASH-NR stained sample from posterior and anterior side; cleared sample imaged from posterior side (the shape shows features of the transparent sample on both the anterior and posterior side; grid: smallest squares 1x1 mm, bold squares 10x10 mm). 3D reconstruction of the entire slice at 16.4 µm isotropic resolution with a Mosaic 16 acquisition (scale bar: 1 cm): The densely stained cell-rich layers are recognizable even at low magnification/large FOV overviews (white arrow heads). The V1/V2 border is indicated by the black arrows in the overview and the enlarged ROI. **b**) Consecutive anterior slice of same occipital lobe approx. 9 mm anterior to occipital pole, panels as described for **a**).

**Figure 4:**
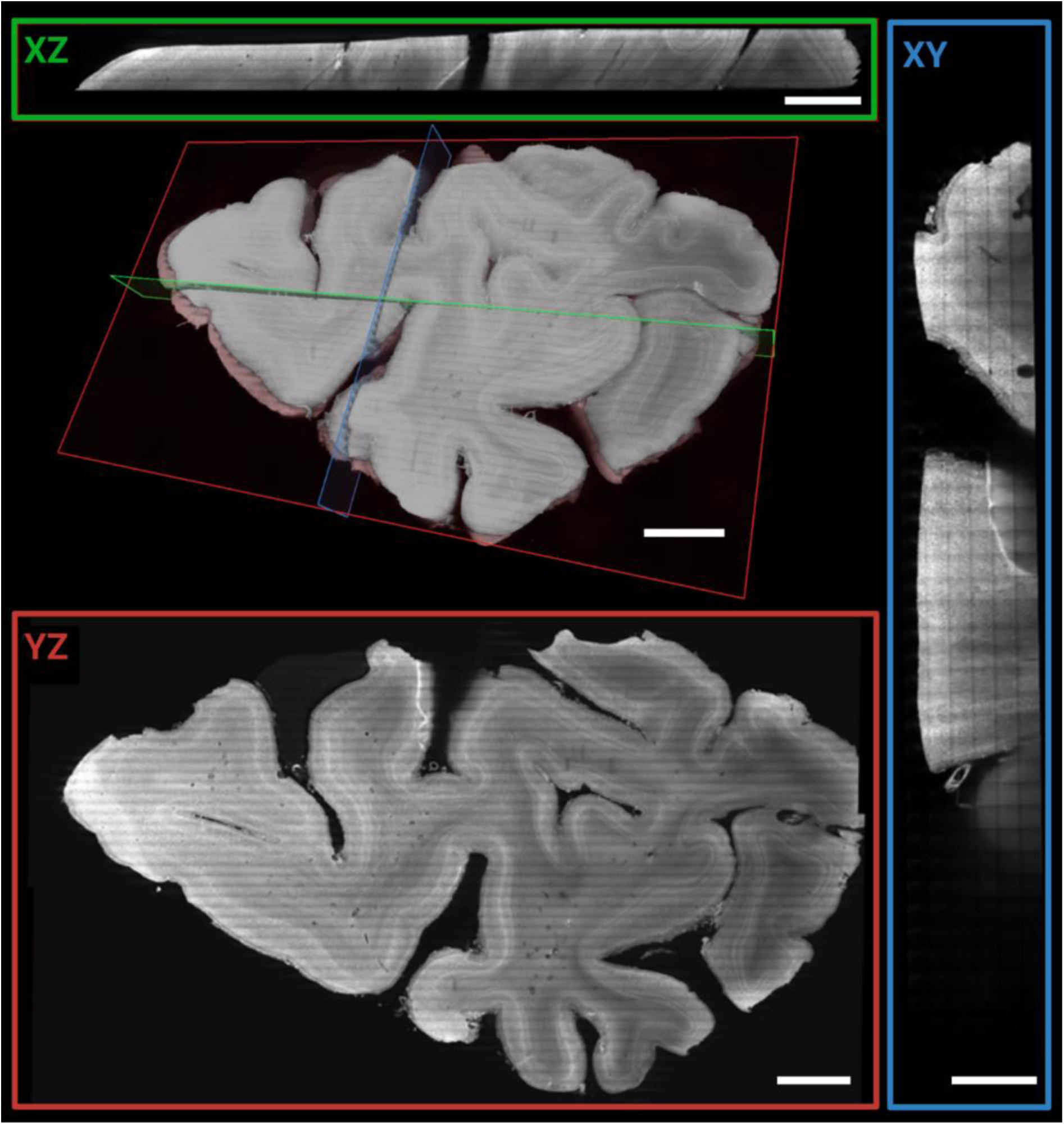
Mosaic 16 overview scan of entire occipital lobe sample. 3D rendering and orthogonal planes of 16.4 µm isotropic data. Orthogonal views of 50 µm MIP (XY: green, XZ: red, YZ: blue) of the entire 3 mm thick sample. Scale bars: 5 mm for XZ, YZ and volume rendering respectively and 2.5 mm for XY plane.

**Figure 5:**
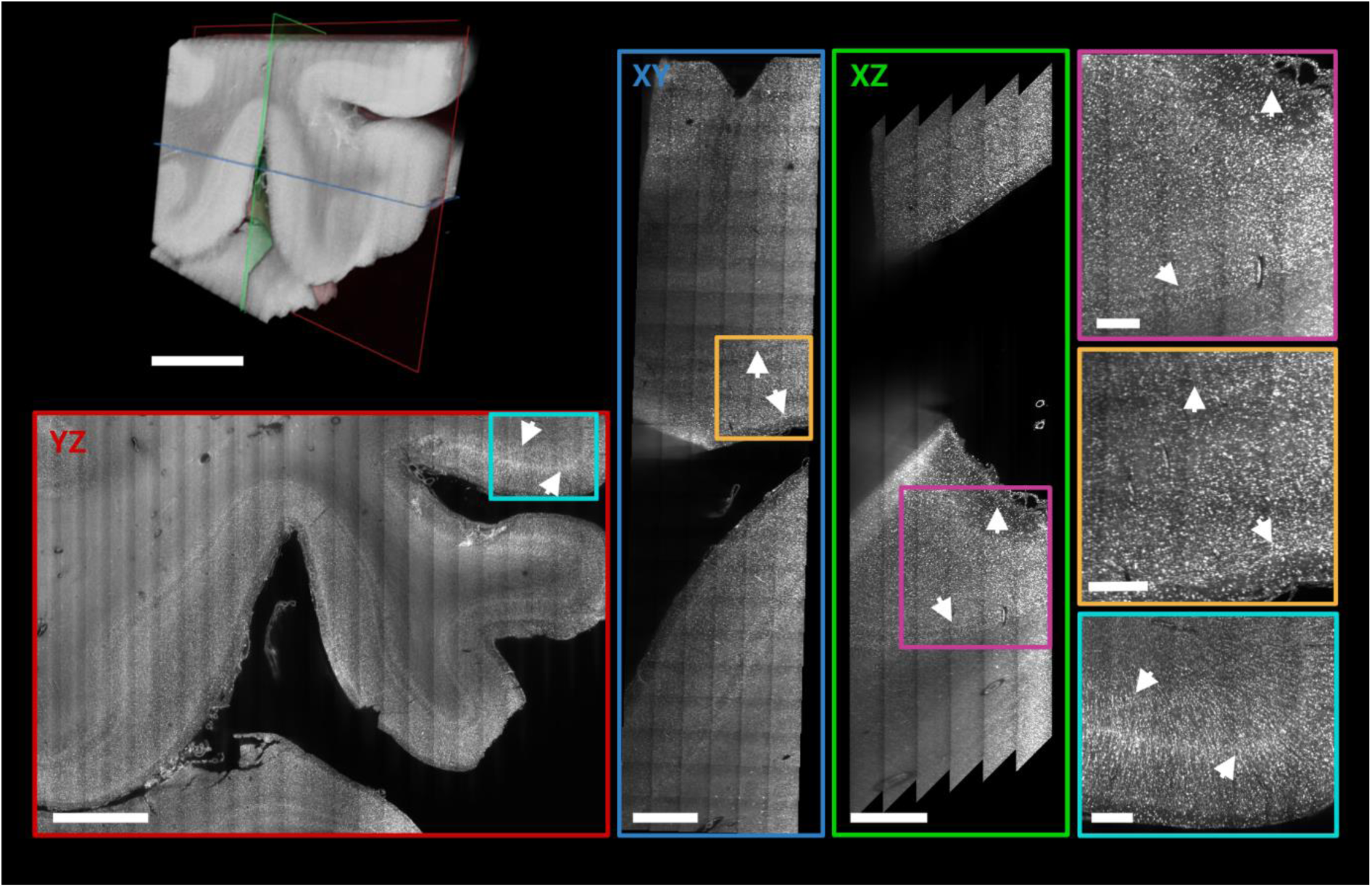
Higher resolution Mosaic 4 scan of human occipital lobe sample. 3D rendering of a Mosaic 4 acquisitions of a ROI in the vicinity of the human primary visual cortex (left top). Orthogonal views of 50 µm MIPs (XY: blue, XZ: green, YZ: red) shows cortical layers independent of the orientation (white arrows). Jagged edges in XZ view result from deskewing when the edge of the acquisition is inside the tissue. Magnified ROIs of the MIP (XY: yellow, XZ: magenta, YZ: cyan) demonstrate qualitatively the isotropically sampled resolution and image and labelling quality deep within the sample. Scale bars: Volume rendering, XY, and YZ plane 3 mm; XZ: 1.5 mm; magnified ROIs 0.5 mm, respectively.

**Figure 6:**
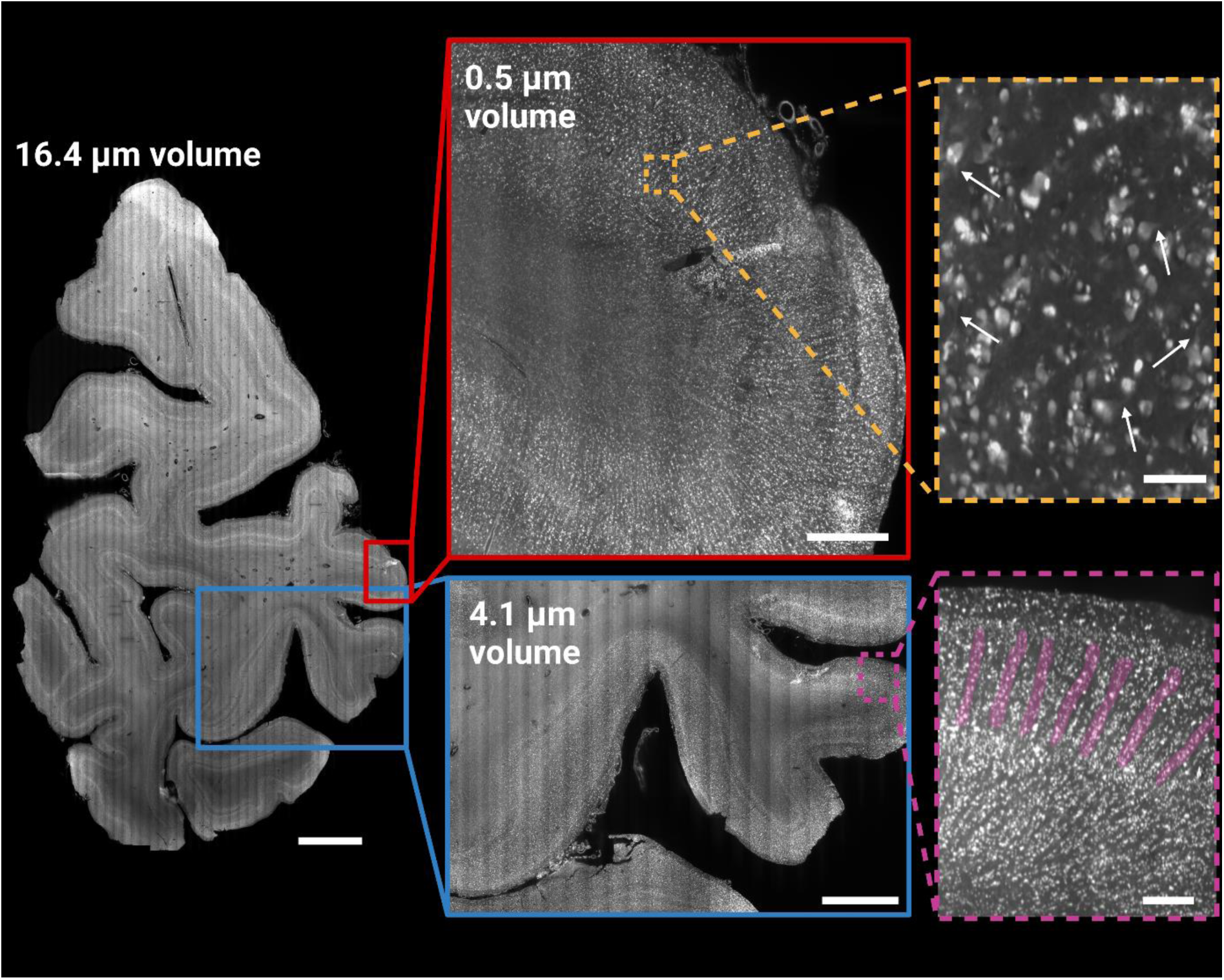
Multi-scale imaging of human brain tissue. Data acquired from the entire occipital lobe slice (third 3 mm thick section from occipital pole towards anterior direction) shown at 16.4 µm resolution (16.4 µm volume; left side). Two gyri were imaged at a resolution of 4.1 µm (blue), and in the 0.5 µm volume (red) at an anisotropic resolution, with a pixel size of 0.725 µm x 0.5127µm x 0.5127 µm. The magnified inserts demonstrate the effective resolution achievable with the 4.1 µm volume (magenta) and 0.5 µm volume (orange), respectively. While the 4.1 µm volume shows mesoscopic structures indicative of minicolumns (highlighted in magenta), it can be difficult to differentiate smaller individual cells from cell clusters at this resolution in the extremely densely populated koniocellular visual cortex. The 0.5 µm volume allows for the identification of single cells, including the proximal parts of apical and basal dendrites (white arrows). Scale bars: 5 mm (16.4 µm volume), 3 mm (4.1 µm volume) and 0.25 mm (4.1 µm volume, enlarged panel), 1 mm (0.5 µm volume) and 0.075 mm (0.5 µm volume, enlarged panel).

**Figure 7:**
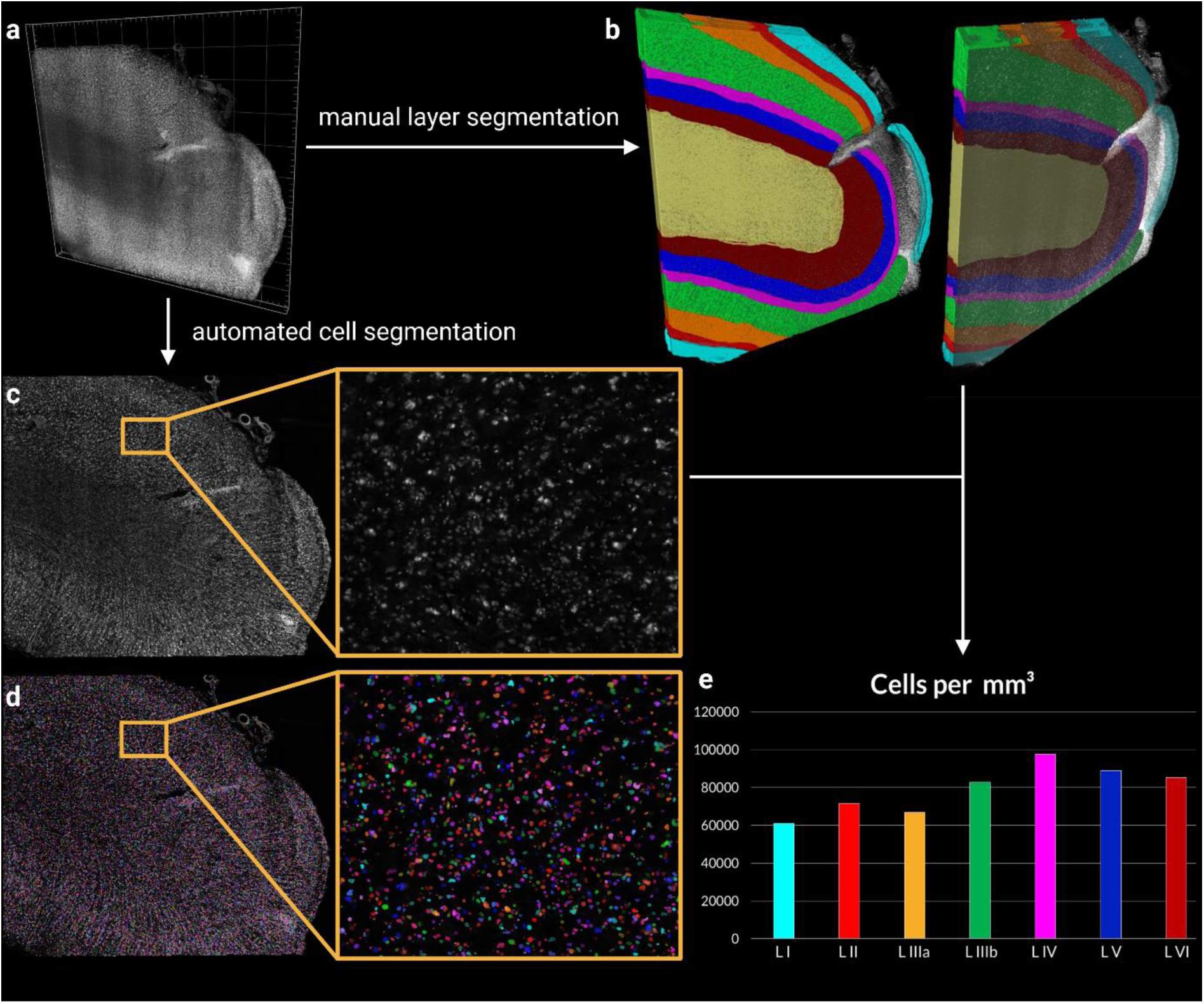
Segmentation and cell counting in a high resolution (0.725 µm x 0.5127µm x 0.5127 µm), single-view volume of human brain tissue. a) Volume rendering of the data set used for the cell counting. b) Results of the manual layer segmentation for the entire volume. Colours are as follows: Layer I cyan, Layer II bright red, Layer IIIa orange, Layer IIIb green, Layer IV magenta, Layer V blue, Layer VI dark red, and white matter in yellow. The tissue in the unsegmented part of the volume was damaged and did not allow for the segmentation of layers and was excluded from the analysis. Single plane (XY plane of the resliced image volume) of the filtered data (c) and automatically segmented objects (d). Segmented cells are shown in random colours. e) Cell density estimates derived from the total count of all the segmented cells per layer segment over the entire image volume (for multiple, unconnected segments of the same layers, the counts were pooled). Colours as indicated for (b).

**Fig. 8:**
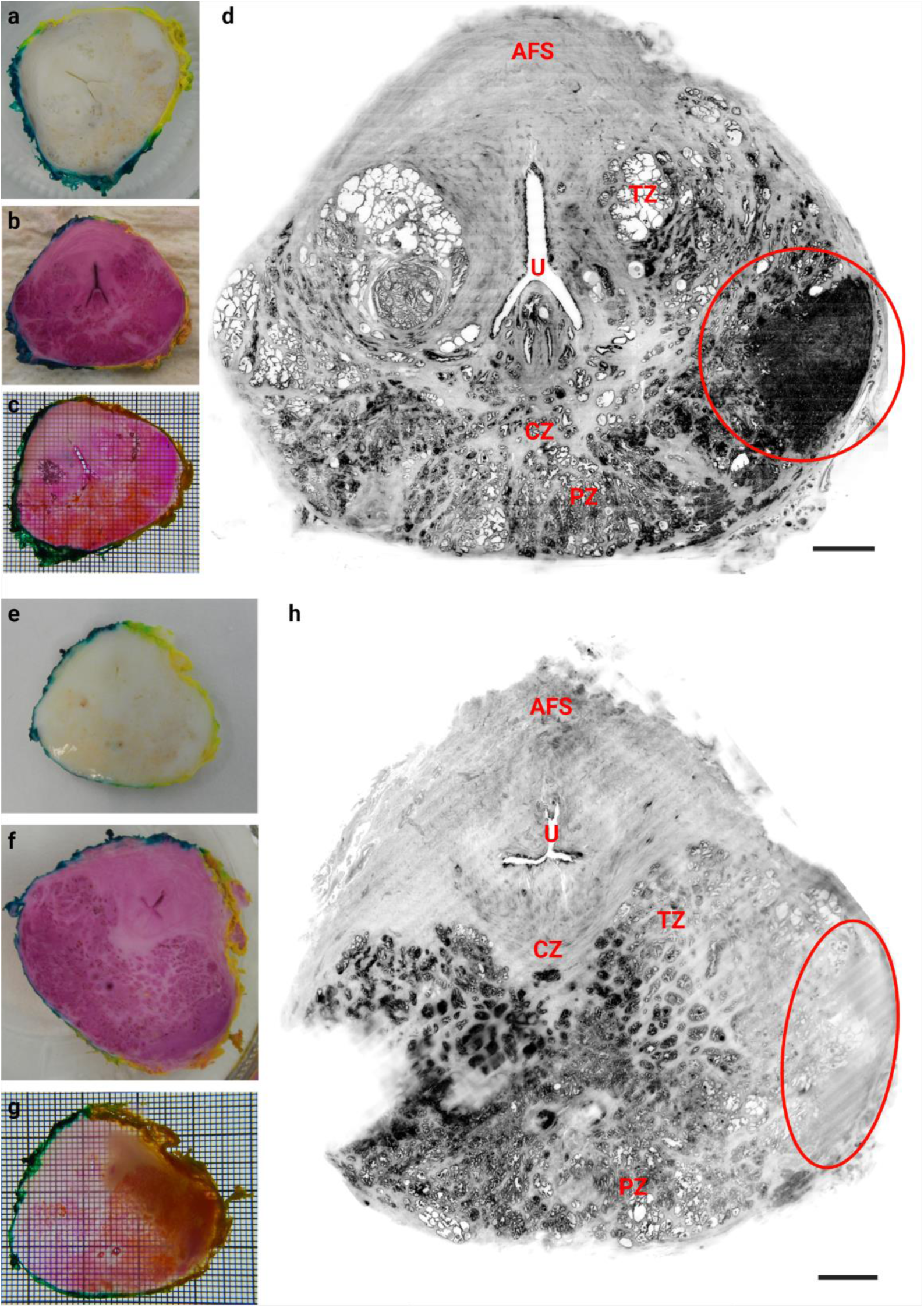
Two prostate cancer samples cleared and labelled with MASH and imaged with ct-dSPIM Mosaic 16. Sample are shown at various stages during the processing pipeline on the left side: After deparaffinisation and bleaching (**a, e**), after staining (**b, f**), and after RI-matching (i.e. cleared, **c, g**). On the right side, a MIPs from the Mosaic 16 data set over approx. 50 µm are shown in inverted greyscale (**d, h**). The red circles indicate the cancerous regions (as identified by a pathologist). Abbreviations: AFS = anterior fibromuscular stroma, CZ = central zone, U = (prostatic) urethra, PZ = peripheral zone, TZ = transition zone. Scale bars: 3 mm.

**Fig. 9:**
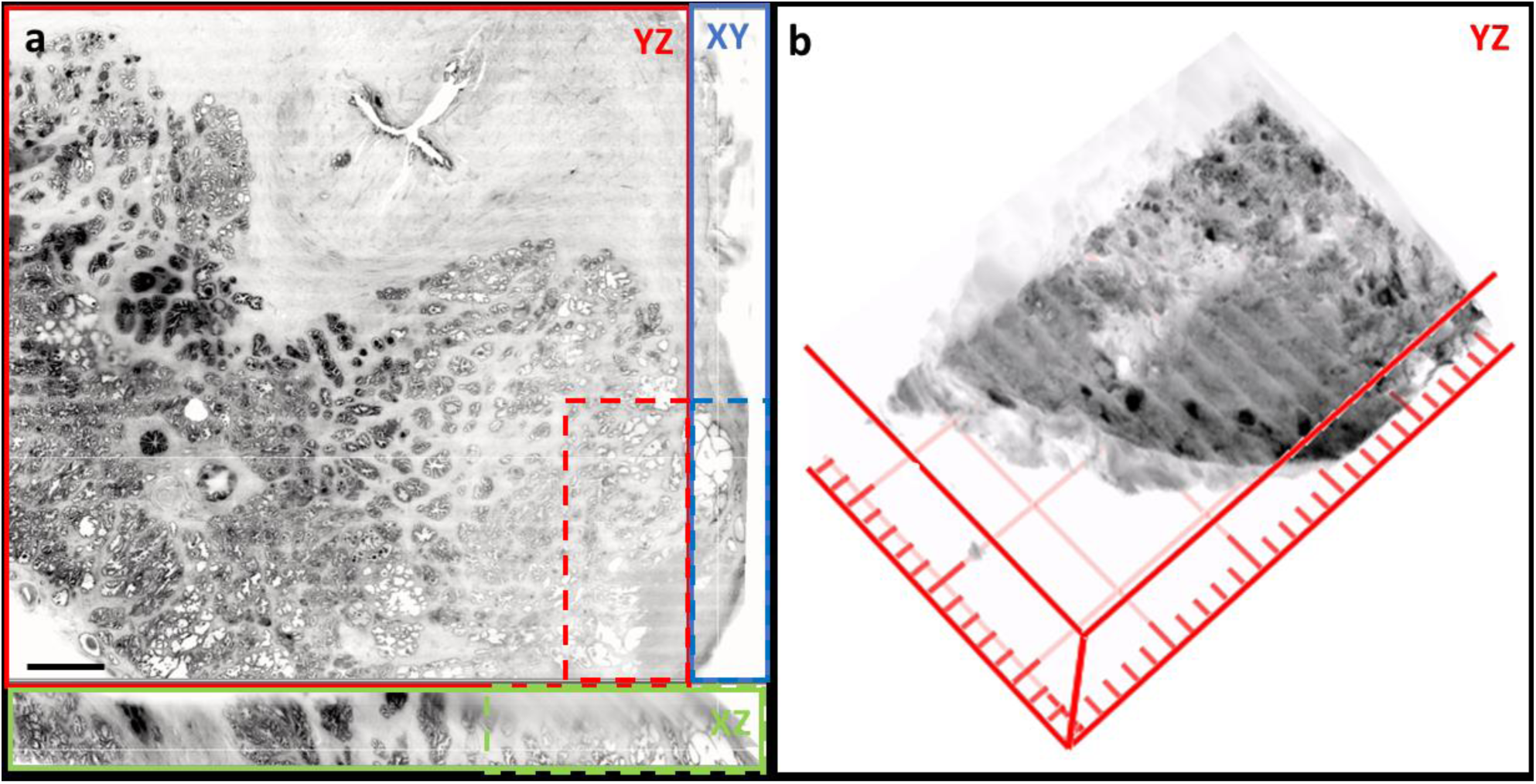
3D rendering and 2D surface view of ct-dSPIM Mosaic 16 prostate cancer sample. Sample from Fig 8 e- h, is shown here in 2D (**a**, one layer across the complete surface in yz shown in the red box, including 2 orthogonal sections across the complete sample thickness in indicated by green (xz) and blue (xy) boxes. Dashed boxes indicate the tumour (as identified by pathologist). The tumour region of this sample (red dashed box in a) is also shown in **b** as a 3D volume rendering. Scale bar: 3 mm.

### Human brain samples

Thick human brain occipital lobe slices (Fig. 3) were taken approx. 6 mm anterior to the occipital pole (third 3 mm thick section in posterior to anterior direction; see Fig. 3 a and Suppl. Video 1) and the consecutive anterior slice (∼ 9 mm anterior to the pole) of same occipital lobe (Fig. 3 b and Suppl. Video 2). The dark discolorations in the unprocessed slices (left-most panels) likely resulted from *post-mortem* accumulation of blood in vessels at the back of the head. Stained slices show the stripe of Gennari in V1 (Fig. 3, indicated by arrows). The morphology of the MASH cleared sample is well preserved after dehydration, delipidation and refractive index (RI) matching, and both grey as well as white matter become highly transparent. The tissue shrinkage observed in MASH cleared samples is relatively minor, with a mean reduction in measured surface are of 12% (suppl. Fig. 2). The black artefacts in Fig. 3 a derive from light-scattering when imaging the hot glue used to fixate the sample in the 3D printed imaging chamber. This occurs when the deepest recorded layer of the image volume extends over the tissue and into the glue below.

We used the occipital lobe samples shown in figure 3 for multiscale ct-dSPIM imaging. First, we performed an overview Mosaic 16 scan of the complete sample (see Fig. 3, 4, 6 and suppl. Video 3). Figure 3 shows the 3D reconstruction of both slices with a Mosaic 16 data volume at 16.4 µm isotropic resolution. The scanning direction can be recognized as the direction of the long imaging stacks or stripes (each 0.74 x 0.74 mm deep and wide and many centimetres long), which are tiled in the remaining two directions to provide full 3D sample coverage. Several cortical layers can be distinguished by their differences in cell density (Fig. 3, white arrow heads) and the V1/V2 border is visible at this mesoscopic resolution as a clear change in layer patterning (Fig. 3, black arrows).

In Figure 4, volume renderings and orthogonal Maximum Intensity Projections (MIPs) are shown to better visualise the volume coverage obtained. The different axes (as labeled in the image viewer, see suppl. Fig. 1) are indicated in green (XZ), red (YZ), and blue (XY) respectively. The XY view shows the tiling of the long acquisition stacks in the volume’s Z direction (corresponding to the X-axis of the stage during the acquisition). The orthogonal views also highlight the label penetration and quality throughout the entire 3 mm thick sample.

After acquiring the overview Mosaic 16, a higher resolution 4.1 µm isotropic resolution Mosaic 4 scan was performed in a ROI close to the V1/V2 border (Fig. 5, 6). The orthogonal views of 50 µm MIPs are indicated by green (XY), red (XZ), and blue (YZ) panels respectively and show differences in cortical layering independent of the orientation (Fig. 5, arrows). The higher magnification insets (XY: yellow, XZ: magenta, YZ: cyan) are a ROI of the MIP and qualitatively demonstrate near isotropic resolution (isotropically sampled at 4.1 µm, but anisotropic optically due to a ∼8 µm thick lightsheet) and high image and labelling quality even deep within the sample (Fig. 5). Mesoarchitectonic cortical features such as minicolums become well visible at this resolution (Fig. 6, magenta panel) and thinner cortical layers, which are not visible in the Mosaic 16 data, can be distinguished. The higher resolution even makes it possible to distinguish some of the largest of individual cells (Fig. 5 and 6, magnified panels), despite the granularity of the occipital cortex.

### Quantitative 3D ct-dSPIM human brain data analysis: Cell counting

To demonstrate the potential of volumetric brain imaging for quantitative 3D histology, we performed automated cell counting in a high resolution 0.5 µm volume (Fig 7; voxel size: 0.725 µm x 0.5127 µm x 0.5127 µm). The aim was to quantitatively compare the cell densities across the different cortical layers. The entire volume (Fig. 7 a) of a deskewed, stitched, and resliced data set was manually segmented into the different cortical layers (Fig. 7 b). The layer segmentation could not be satisfactorily performed in one region of the data set, which contained damaged tissue. This region was therefore excluded from the analysis (uncolloured parts in Fig. 7 b). Subsequently, the data was processed in an automated pipeline to filter the raw images and segment cell bodies (Fig. 7 c and d). The number of all cell segments, larger than 125 µm^3^ and fully contained within the corresponding layer segment, was used to derive cell density estimates for each layer (Fig. 7 e). The cell counts were pooled for layers with multiple unconnected segments. As expected, layer I shows the lowest cell density with just over 60000 segmented objects per mm^3^, although the density is higher than expected for this layer. Layer II, while showing higher cell densities than layer I or IIIa (as expected), shows a lower density than expected from the qualitative impression during layer segmentation. Surpisingly, layer IIIb, V, and VI had higher cell densities than layer II or IIIa. Layer IV, which is visibly by far the densest layer, also exhibited the highest cell density in our data set, with almost 100000 segmented objects per mm^3^. Several of these observation are likely at least partially explained by partial volume effects. To validate the automated segmentation results, we performed manual cell counting on the same data set. The employed automatic cell segmentation method employed to an average overcounting of 36%, compared to the manual counts (see Suppl. Fig. 3). This was partially, but not entirely alleviated by the exclusion of segments that were considered too small to represent cells (< 125 µm^3^). After filtering these small structures, the second automated results overcounted by an average of 17%. It was observed that especially larger cells in the automatic segmentation were often segmented into multiple smaller objects when there were visible intensity differences across the cytoplasm or between cytoplasms nucleus, and nucleolus.

### Axial whole-mount prostate sections

We developed FFPE-MASH for formalin-fixed and paraffin-embedded (FFPE) human axial whole-mount prostate sections. For the first time the MASH protocol with the MASH-NR label has been successfully applied to large prostate cancer samples from different patients (the axial whole-mount sections; Fig. 8, 9; suppl. fig. 4, 5; suppl. video 5). We therefore demonstrate the feasibility for the application of MASH not only to other-than-brain human organ tissue, but also on FFPE material. Moreover, we were able to show the feasibility of high-speed (1.7h/1cm^3^) Mosaic 16 ct-dSPIM imaging on large MASH-NR cleared and labelled prostate cancer samples. The large mesoscopic overviews allow for the anatomical description of prostate tissue morphology and the indication of possible neoplastic regions (Fig. 8, d, h; indicated by red circles), which was confirmed to be a prostate adenocarcinoma by histopathology. Microscopic evaluation of the corresponding haematoxylin-eosin sections showed that the tumour consists of cribriform and fused neoplastic glands, compatible with a high-grade prostate adenocarcinoma. Different layer zones and compartments of the prostate gland could be classified in the Mosaic 16 volume (Fig. 8 d, h), namely the fibro muscular stroma (AFS), the central zone (CZ), the peripheral zone (PZ), the transition zone (TZ) and the urethra (U). Furthermore, higher resolution Mosaic 4 scans allowed for the detection of the tumour morphology throughout the tissue sample and the pathologist suggested a Gleason Score of both 3 and 4 in the axial whole-mount (prostatectomy) section (suppl. Fig. 5).

### 3D visualization of prostate ct-dSPIM data

The 3D visualization of the ct-dSPIM imaged the axial whole-mount prostate sections (Fig. 9; suppl. fig. 5) allows for the detection of the cross-cut of the urethra prostatica, shown in the orthogonal sections across the complete sample thickness (Fig. 9 a, green and blue box, dashed insert indicates the tumor region). The 3D volume shown in Fig. 9 also shows the cancerous region, indicated with the red dashed line in a in the surface view, as confirmed by a pathologist. The tumor regions are also indicated in the orthogonal section in Fig. 9a in the green and the blue dashed line. Fig. 9b shows a 3D higher maginification of the tumor region shown in the red dashed line box in a.

## Discussion

### Novel Mosaic Scans of large cleared human tissue samples with ct-dSPIM prototype

Here we present the novel ct-dSPIM prototype for cleared tissue light-sheet imaging of large-scale human tissue samples and our application to both human brain and prostate axial whole-mount sections (after prostatectomy). We were able to demonstrate the efficiency and feasibility of ct-dSPIM imaging with both MASH prepared [23] archival human occipital lobe (Fig. 3-7) and prostate resections (prostatectomy) (Fig. 8, 9). The MASH protocol [23] with the MASH-NR labelling was successfully applied to large formalin-fixed and paraffin-embedded (FFPE) prostate cancer samples for the first time, as shown in figures 8 and 9, as well as in suppl. figures 4 and 5 and suppl. movie 5. This does not only show the applicability of MASH to other tissue types besides human brain but, equally important, to FFPE samples in addition to formalin-fixed samples. Previously, we applied the MASH protocol to formalin-fixed archival human brain samples that were kept in 4% PFA until use. The use of the MASH protocol on FFPE material could increase its application in other domains, since FFPE tissue has been commonly used in research and clinical application for decades, which means the potential effect on clinical practice is considerable. The most important step in the modification of MASH towards FFPE tissue is the initial deparaffinisation in xylene, which must be sufficiently long to allow the complete dissolution of the paraffin in thick samples. Thus, this step might be omitted by application of this technique on fresh samples, without prior fixations and processing with paraffinisation, which would considerably speed up the MASH tissue processing time.

Complimentary to overview Mosaic 16 acquisitions with an isotropic resolution of 16.4 µm, we have acquired higher resolutions scans within specific regions of interest. Whereas a Mosaic 16 provides a whole tissue block overview, Mosaic 4 (4.1 µm isotropic resolution) and Mosaic 0.5 (near 1 µm resolution) allow for a more detailed assessment of parts of the same tissue. Together, this multiscale data acquisition scheme allows for the identification of different mesoscopic features, such as layers, minicolumns, and cell bodies in the brain (Fig. 6). This enables the comprehensive and detailed analysis of large human tissue samples.

The isotropic sampling of the Mosaic 16 and Mosaic 4 data sets is based on a √2 relation of slice step length to lateral sampling resolution and is optically limited by axial resolution, i.e., by the theoretical ∼8 µm light-sheet thickness. The overall acquisition speed of Mosaic scans is only limited by the maximum stage scanning-speed and currently ranges from 1.7h/∼1 cm^3^ for Mosaic 16 to 5h/∼1 cm^3^ for Mosaic 4 and 15.8h/∼1 cm^3^ for Mosaic 0.5. Mosaic 16 overview scans allowed for high-volume high-speed visualisation of entire 3mm prostatectomy tissue slab or an entire 3mm human occipital lobe tissue slab. Here, we show Mosaic scans of the occipital samples with an acquisition duration between 2 hrs 23 min (Mosaic 0.5), 3 hrs 45 min (Mosaic 4) and up to 8 hrs 26 min of the entire 5cm^3^ large occlobe sample (Mosaic 16; Fig. 3-7). The duration to acquire one surface layer Mosaic 16 scan of one prostate sample, as shown in Fig. 8, was 50 min.

In the future, the method could push current limits on resolution. It is theoretically possible to perform ∼1 µm effective optical resolution dual view ct-dSPIM imaging, which has mostly been demonstrated in cell cultures. Very recently, demonstrations in cleared tissue have emerged [20] [29] applying dual view imaging and subsequent dual view deconvolution processing through strong speed-ups of the deconvolution processing. However, even in the most recent developments [29], where the deconvolution is implemented through deep learning informed by the image formation process, processing times are still on the order of a few hours (2-3h) for volumes three orders of magnitude smaller (1.4x2.3x0.5mm3) than the ones reported here, which implies thousands of hours of processing for the very large samples presented here with even the most advanced current methods. Furthermore, dual-view acquisitions would double the current acquisition times, making the total time cost for the resolution improvement considerable and unrealistic in the relevant context of very large tissue samples and future clinical application in the prostate. Moreover, for the purposes of this study, with an aim at efficient and scalable large FoV imaging at sufficient resolution (rather than the highest possible optically limited resolution), the cost/benefit analysis and feasibility in terms of total time makes dual view imaging untenable. In our current study we have included ct-dSPIM mosaic imaging data of a human brain (occipital lobe) sub volume with a (sampled) resolution of 0.725 µm x 0.5127µm x 0.5127 µm. Although the true optical resolution is lower (on the order of 8.0µm x 0.8µm x 0.8µm with the 0.4 NA objective), these single view data can be acquired at reasonable data rates and are sufficient for the intended purposes including single cell detection, segmentation and quantitative cell counts in human brain samples. In prostate cancer samples a sampled resolution ranging from 16-0.5 µm enables efficient 3D tumour imaging at an image quality sufficient for clinical investigation (Fig. 9,8; suppl. Fig. 4,5). For future work we see opportunities (e.g. subcellular resolution for intra-cellular structures or neuron axonal or dendritic structures) and possibility to combine our current efforts with higher near-isotropic resolution. In such efforts, both dual view deconvolved approaches and single view axially scanned approaches will deserve consideration for large sample brain and prostate imaging [30, 31].

### Human brain cell segmentation and counting

One of the promising aspects of volumetric imaging shown in this paper, is the fact that it enables us to image a large volume of human cortex at a high spatial resolution, which in turn allows us to investigate cellular structures over extended fields of views. In the future, this could in principle allow quantifying cell densities for different areas and layers without stereological bias, since the derived cell counts are not extrapolations from 2D sections. However, at this point, stereologically derived estimates of cell densities should still be considered the gold standard as they are well established. It is, hence, interesting how our 3D segmentation and counting results relate to stereological data in the literature. Unfortunately, cell density estimates in the human secondary visual cortex are rare. Estimates that have been calculated with stereological methods, such as the isotropic fractionator [32], are typically given in cells per gram of brain tissue and are not straightforward to convert to our volume estimates.

Leuba and Garey [33] found average cell numbers in area V2 of 147600 under a cortical area of 1 mm^2^, or 63700 cells/mm^3^. The total cell density in our data set across all layers was 81903 cells/mm^3^. It is considered unlikely that these higher cell counts in our data arise from clearing induced tissue shrinkage, as the observed change in size of our samples is relatively minor (see suppl. Fig. 2). This is especially true when compared to other solvent-based tissue clearing methods, which show much higher tissue shrinkage of 30% [14] or even up to 50% [34]. If we take the observed overcounting of the automated cell segmentation of, on average, 17% into account (Suppl. Fig. 3), our corrected cell counts of 67980 cells/mm^3^ are relatively close to Leuba and Garey. When focussing on the individual layers, our cell density in layer IV is generally in good agreement with Leuba and Garey with 97529 cells/mm^3^ (our data) vs 107316 cells/mm^3^[33]. To derive densities of all cells, we multiplied their reported neuron numbers with their reported neuron/glia ratio and added these two populations, as they only provide whole cell population densities for the entire area, but not in their layer-specific table. Unfortunately, Leuba and Garey do not list densities for all layers, but report supragranular layers II and III, as well as infragranular layers V and VI together. They report a supragranular cell density of 63558 cells/mm3, which is similar to the number of cells we found in layer IIIa (66884 cells/mm^3^). However, both layer II (71599 cells/mm^3^) and IIIb (82712 cells/mm^3^) show higher cell numbers and, therefore, the combined supragranular cell density is higher as well, with 77807 cells/mm^3^. As mentioned earlier, the very high cell density observed in layer IIIb was surprising, given the visual appearance of that layer as less dense with sparser, larger pyramidal neurons. A possible explanation for this discrepancy could be a partial volume effect from the very dense adjacent layer IV, arising from imperfect manual layer segmentation. Another likely contributing factor could be an observed tendency of the automated cell segmentation to split larger neurons into multiple segments when they show intensity differences over their cytoplasmic volume (see Suppl. Fig. 3). This tendency to segment larger cells into multiple segments led to an average overcounting of the automated cell segmentation of about 17%, after segments smaller than 125 µm^3^ had been filtered out. If we take this overcounting into consideration, the combined supragranular cell count of 64580 cells/mm^3^ is considerably closer to Leuba and Garey. Both of these factors might also explain the larger cell densities observed in our data for layers V, since the high density of layer V (88781 cells/mm^3^) was similarly unexpected. Partial volume effects could not explain the similarly high density of layer VI (81904 cells/mm^3^), although the splitting of large cell bodies likely still lead to an overcount. Hence, our combined infragranular cell density is substantially larger than one reported by Leuba and Garey (86579 cells/mm^3^ vs 53658 cells/mm^3^). Even if we account for the overcounting of the automated segmentation, our observed infragranular cell densities would be higher than that of Leuba and Garey (71861 cells/mm^3^). Improvements in the segmentation quality could alleviate at least some of these discrepancies and bring cell estimates in this area closer to the stereological estimates. Zhao et al. use a deep-learning-based segmentation on volumetric data of cleared human brain [14]. It should be noted that the authors of this paper report much higher densities (162000-216000 cells/mm^3^) than either Leuba and Garey or the work reported here, although they did not investigate different areas.

One possible aspect that could account for varying cell numbers across these approaches, is also the amount of shrinkage introduced during the histological processing. Since the tissue clearing approach used in this study is different from Zhao et al. [35], and known to produce relatively minor shrinkage (12% compared to 30% reported in Zhao et al.; see Suppl. Fig. 2), this could account, at least in part, for this discrepancy. Another recently published study that applied a very sophisticated cell segmentation pipeline [36] came again to vastly different results with neuron densities of 14000- 24000 neurons/mm^3^ (which means around 28000-47500 cells/mm^3^ when applying Leuba and Garey’s neuron/glia ratio). The clearing approach used in this study is an aqueous tissue clearing protocol, which expands the tissue. This in turn might explain the lower estimates. In addition, they rely on antibody labelling to identify neuronal cell populations, rather than a more general organic dye with smaller molecular weight, which could also account for some of the observed differences. Lastly, it is also possible that the stereological 2D method deployed by Leuba and Garey leads to systematic undercounting, when extrapolating the 2D cell counts on sections to a large volume of tissue.

### Tumour and tissue structure visualization of prostate resection sections with the ct-dSPIM

Classical histo-pathological prostate cancer diagnostics, in particular of multifocal tumours is challenging [3]. Moreover, the assessment of the entire core biopsy, or the resection of the entire sample would take up to 10 days, which is inefficient and very expensive, and almost never performed in standard-of-care clinical practice. Therefore, possible Mosaic ct-dSPIM imaging of prostate, in collaboration with oncologists and pathologists at the local hospital open up new avenues for future cancer diagnostics.

Recent work has demonstrated the usefulness of LSFM to examine 1- 2.5 mm thick human prostate core-needle biopsies by means of a home-built open-top LSFM system [37]. Instead, here complete 5 mm thick axial whole-mount prostate sections were examined with both Mosaic 16 and 4 ct-dSPIM scans within approx. 1-5 hours, which is time efficient and enables a relatively fast overview assessment of the entire biopsy (Fig. 8, 9; suppl. Fig. 4,5; suppl. Video 5). Additionally, this allows for a retrospective examination of a very high number of archival prostate cancer samples (up to 8 samples per day). High-volume 3D imaging can enable the visualization of tumours in deeper layers or regions, and throughout the entire thick prostate resection sample. This has the potential to provide new insights into both cancerous, as well as benign, prostate tissue structure. To date the current information on 3D prostate architecture is limited, as it is based on MRI examination, which does not allow a detailed and high-resolution analysis of the complete organ, including gland formation, lumen and epithelial layer. However, in the current histo-pathological practice, the most frequently used technique is a combination of classic bright-field microscopy and three to four levels of 5 µm thick tissue sections, resulting in 2D tissue information. This can lead to under grading of the tumour or false-negative diagnosis in case of the multifocal carcinoma, because these are mostly present on the deep levels of tissue blocks. Therefore, it is of high value for the pathologist and oncologist to get novel 2D and 3D high-resolution high-volume insight into detailed prostate tumour architecture using 5 mm thick patient prostate samples (Fig. 8, 9; suppl. Fig. 4, 5; suppl. video 5). Quantitative light-sheet imaging of those thicker prostate cancer samples (prostatectomy) can improve the precision of the diagnostic. The results of this study show that it is possible to detect tumours with our method of cleared tissue ct-dSPIM imaging and even enable Gleason score grading (suppl. Fig. 5), including the detection of the neoplastic process and to distinguish normal benign prostatic tissue from adenocarcinoma. Both Mosaic 16 (Fig. 8, 9) and Mosaic 4 scans (suppl. Fig. 4,5; suppl. Video 5) allowed for the detection of tumours in the axial whole-mount prostate section from different patients.

Moreover, since Mosaic acquisitions on the ct-dSPIM allow for high-speed 3D examination of large-scale prostate samples, it is feasible to develop high-throughput imaging pipelines, for approx. 20 samples in 10 days, if only one sample is imaged at a time. The standard processing of prostate includes at least 24 hours of fixation, followed by grossing and processing in a vacuum infiltration tissue processor (at least 24 hours). Thus, this step might be omitted by application of the proposed MASH and ct-dSPIM approach on fresh resection specimen. This could potentially have a considerable effect on the clinical practice and accelerate diagnostic workflow. Given the size of the imaging container (see Fig. 1), multiple samples could be accommodated in parallel in the ct-dSPIM, potentially making the imaging yet more efficient. This can result in rapid gathering of knowledge and is feasible for various patient samples at different stages of the disease while providing sufficient data for statistics and quantification of tumour variation and localization.

In summary, our method has the potential to improve the imaging of human prostate cancer resections samples. It provides novel 3D visualization of the entire sample in detail, which is infeasible in time and resources with the traditional methods. It could also lead to faster prostate imaging, which has benefits for the efficiency of clinical practice, as well as lead to new opportunities for (archival tissue) research. Additionally, it could provide more accurate diagnostics, especially in multi-focal carcinomas.

### Future perspectives

We demonstrated large scale Mosaic imaging on the ct-dSPIM system and showed that we were able to overcome limitations of other existing light-sheet microscopes for very large human sample coverage [17, 38]. The ct-dSPIM uses an oblique geometry with the objectives dipping directly into the imaging liquid from above, unlike other recently published set-ups [38, 39]. In earlier systems, the objectives are located beneath the sample, which necessitates imaging through the glass bottom of the chamber. The sample size is potentially further limited by this geometry as the working distance (WD) is effectively lost by imaging through the glass and an additional lens. Other systems use a more standard (i.e. upright, not oblique) light sheet geometry and use a novel way of generating the light sheet itself to image very large samples [11]. In these systems, the lateral extent will ultimately be limited, as even in the most transparent samples, light scattering will occur to some degree and the image quality deeper into the tissue deteriorates. This can be avoided with an oblique set-up such as the ct-dSPIM, leaving the lateral size of the sample limited only by the travel range of the microscope stage.

Further hardware changes, such as a cylindrical lens (rather than digital scanned laser light-sheet) scanner, and stage tiling trajectory adjustments, such as a serpentine trajectory, could potentially further increase imaging speed 2- to 4-fold. This would open up a completely new scale in the investigation of *post-mortem* healthy and diseased brain tissue. Other hardware modifications of the ct-dSPIM set-up, such as higher NA and magnification objectives, as well as an axially scanned light sheet approach, could potentially allow for an increase in resolution for imaging at the cost of field of view or time to image the same sample size. These future developments might allow imaging different structures and markers in larger parts of the brain, with the potential to provide novel insights into healthy and pathological human neuroanatomy.

Moreover, establishing further labelling options, such as small molecule labelling of brain angio- architecture, as well as antibody labelling strategies which are also suitable for ct-dSPIM large sample imaging, could further extend possibilities. In the future, the entire pipeline from tissue processing (with MASH clearing and labelling) to ct-dSPIM Mosaic imaging and data analysis could be extended to larger parts of the brain and brain regions, such as the whole temporal or occipital lobe in 3-5 mm thick slabs. As the lateral sample size can potentially be much larger in the current set-up, it would even be possible to image whole brain slices, provided a tissue-processing pipeline for samples of this size is established.

Although we demonstrated the use of MASH prepared human and prostate as application cases for large scale acquisitions on the ct-dSPIM, this technique could potentially be extended to a variety of other human and non-human mammalian tissues as well. This could make the combination of MASH with ct-dSPIM imaging a powerful tool for anatomical and pathological studies in general.

## Acknowledgements

We thank Maurizio Abbate (Arivis AG) for kindly providing the custom-made python script for the Vision4D environment to divide the data volume was divided into a 100 x 100 µm grid. Moreover, we thank Dr. Jon Daniels (Applied Scientific Instrumentation) for technical advice and providing Fig. 2 with the optical layout of the ct-dSPIM.

## Appendix

**Supplementary Video 1.** Plane-by-plane view of the resliced Mosaic 16 scan of occipital lobe slice 1. The sample is viewed along the YZ plane, to visualise its full extend. The volume is represented in inverted greyscale.

**Supplementary Video 2.** Plane-by-plane view of the resliced Mosaic 16 scan of occipital lobe slice 2. The sample is viewed along the YZ plane, to visualise its full extend. The volume is represented in inverted greyscale.

**Supplementary Video 3.** 3D rendering of an entire human occipital lobe slice 2 (3 mm thick). The slice was imaged with a Mosaic 16 scan. The mesoscopic and isotropic resolution of the dataset is sufficient to appreciate large anatomical landmarks and differences in the cytoarchitecture.

**Supplementary Video 4**. 3D rendering of the single view volume (human occiptital lobe) obtained at the highest resolution possible with the ct-dSPIM. ROIs (red spheres) were places manually along the volume, which was rotated and clipped to evaluate the placement of the ROIs within the layers.

**Supplementary Video 5:** Changes of prostate morphology in 3D. The video shows an average intensity projection of the prostatectomy slice (left, second 1-2), with normal prostatic glands of the transition zone highlighted in cyan and the tumour region highlighted in yellow (right), respectively. Second 4-17 show the change of prostate morphology as one moves plane-by-plane through the resliced data set. This allows to trace the changing shapes of both benign, as well as cancerous tissue elements.

---

**Supplementary figure 1:**
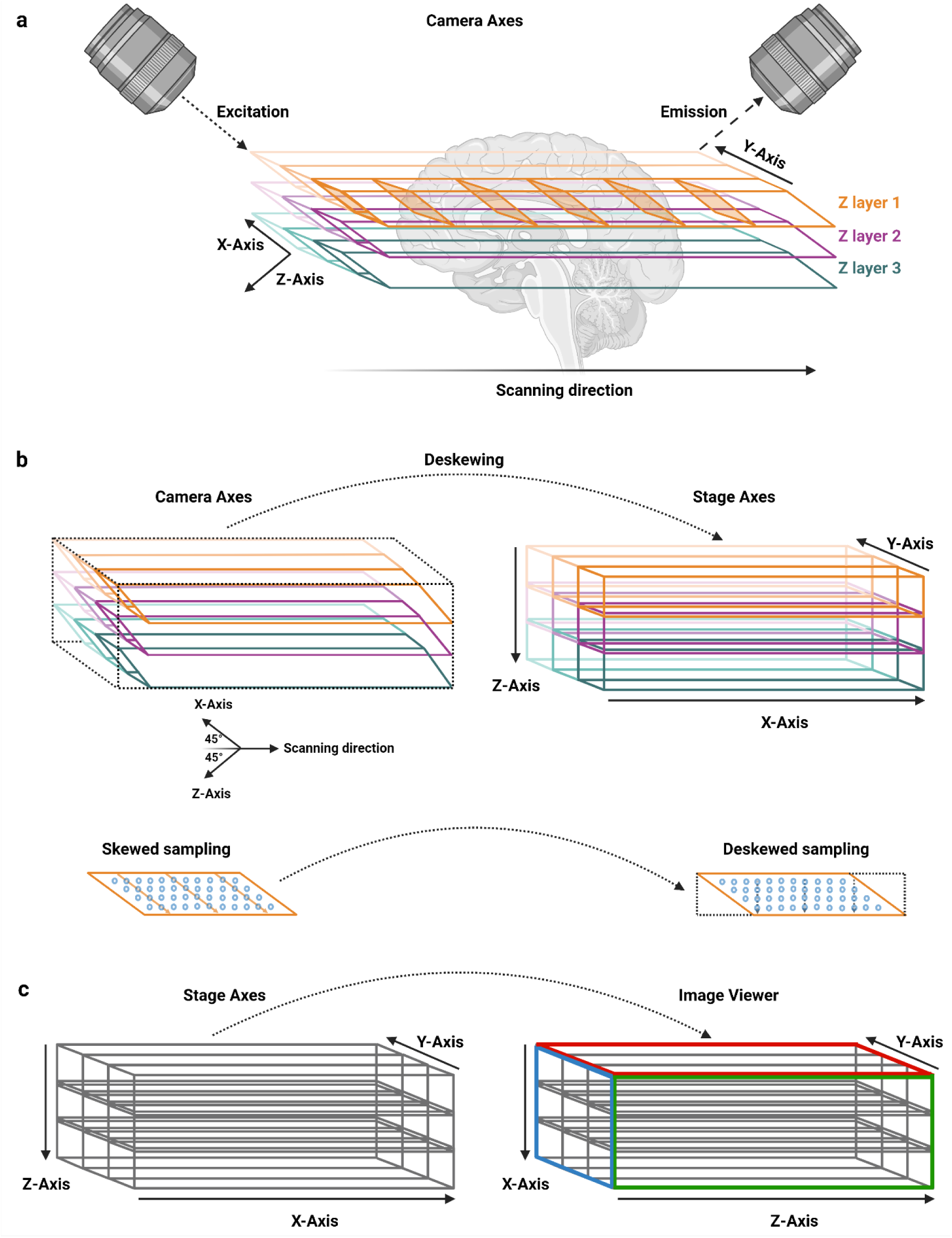
Camera and stage axes in ct-dSPIM imaging, and image viewer axes and their relationships. a) When acquiring image volumes with the ct-dSPIM, which uses stage scanning to move the sample through the imaging plane (i.e. the light sheet) the axes do not correspond to orthogonal XYZ coordinates. Instead, the camera Z-axis (perpendicular to the imaging plane) is at a 45° angle towards the scanning direction which is performed along the x-axis of the stage. Similarly, the camera X-axis is at a 45° angle towards the scanning direction. The Y-axis is the only axis that is shared between camera axes and stage axes. Therefore, stage-scanned image acquisition leads to a skewed parallelepipedal stack shape with non-orthogonal axes, which will be warped when viewed in the orthogonal axis system b) Shows how the deskewing operation transforms the skewed parallelepipedal stack (top, left), into a rectangular stack (top, right). If the stage step size along the scanning direction is chosen equivalent to the final, downsampled pixel size of a Mosaic acquisition (and therefore has a sqrt(2) relationship to the in-plane XY resolution in camera axes), the deskewing operation is equivalent a fast resampling of the skewed data, without the need for slow interpolation (bottom), although interpolation can be performed for every chosen stage step size c) As is the case for most 3D imaging data, image viewer software may use its own axis labelling which may be different from the that of the data imported into the viewer. The image viewer axis labeling here is the one used in figures in this work. Note that the deskewing step in b and axis relabeling step in c may be combined into the same processing step or reversed in order in actual software implementations.

**Supplementary figure 2:**
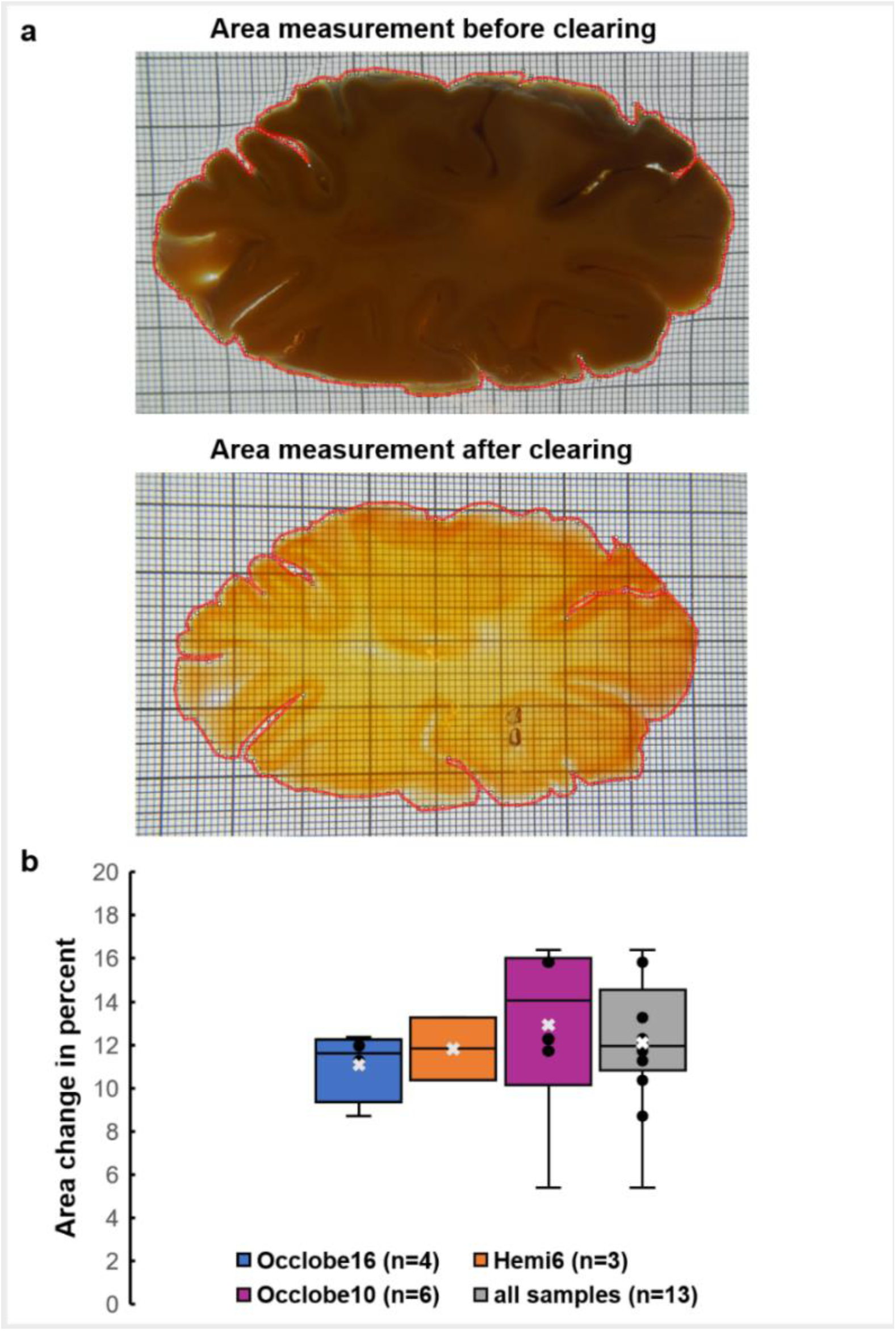
Tissue shrinkage after the MASH clearing protocol. a) Example of the surface area measurement before (top) and after (bottom) MASH clearing. b) Shows box-and-whisker plots of the area change in percent for all 3 donors, as well as the pooled data from all 13 brain slices. Whiskers and box compartments show lower and upper quartiles, and interquartile range, respectively. The black horizontal line indicates the median and the white cross the mean of the data. Individual data points are shown as black dots, unless they would be located directly on the box or whisker lines.

**Supplementary figure 3:**
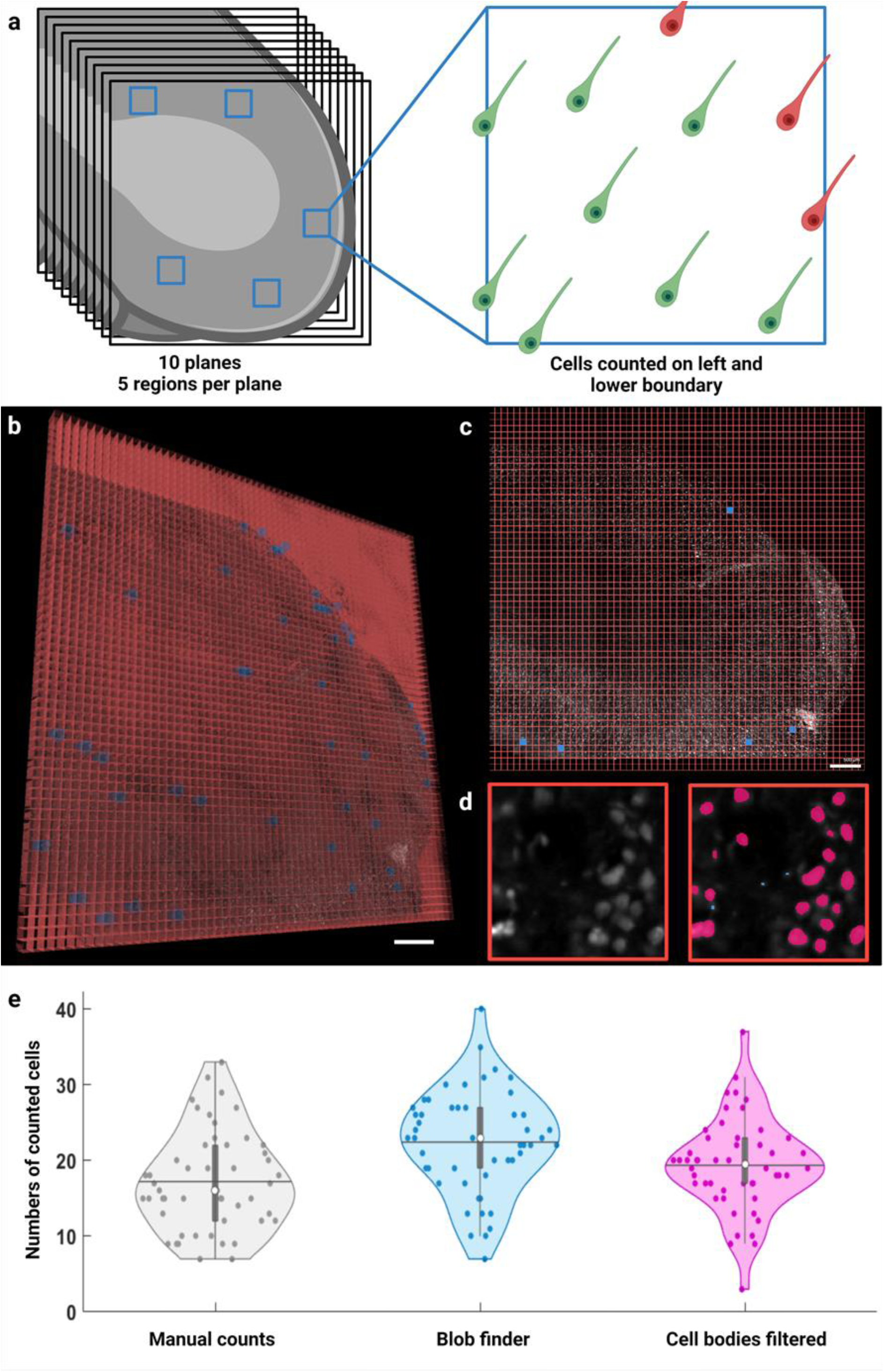
Manual validation of automated cell segmentation. a) Schematic representation of the validation procedure. 10 planes from the same dataset as shown in figure 7 were pseudo-randomly selected using a random number generator. On each plane, 5 ROIs (100x100 µm), selected in the same manner, were used for cell counting. Cells were counted that were either fully contained within the ROI or intersecting with the left and lower boundary. b) 3D rendering of the entire dataset, with the 100x100 µm grid overlaid in red. All ROIs that were counted on all 10 planes are depicted in blue. c) One representative plane out of the 10 selected ones, with the grid overlaid in red and the 5 ROIs displayed in blue. d) Single ROI showing the raw data (left) and the automatic segmentation results. The filtered segments, taking only object larger than 125 µm3 into account, are shown in magenta. The smaller segments which were detected by the “blob finder” segmentation of Vision4D, but have been filtered out, are shown in light blue. e) Violin plots for the manually counted cells (grey), the unfiltered segmentation results (“blob finder”, blue) and the filtered segmentation results (“cell bodies filtered”, magenta). White dots within the box plots indicate the median, the horizontal lines represent the mean for each dataset. Scale bars: 500 µm each.

**Supplementary figure 4:**
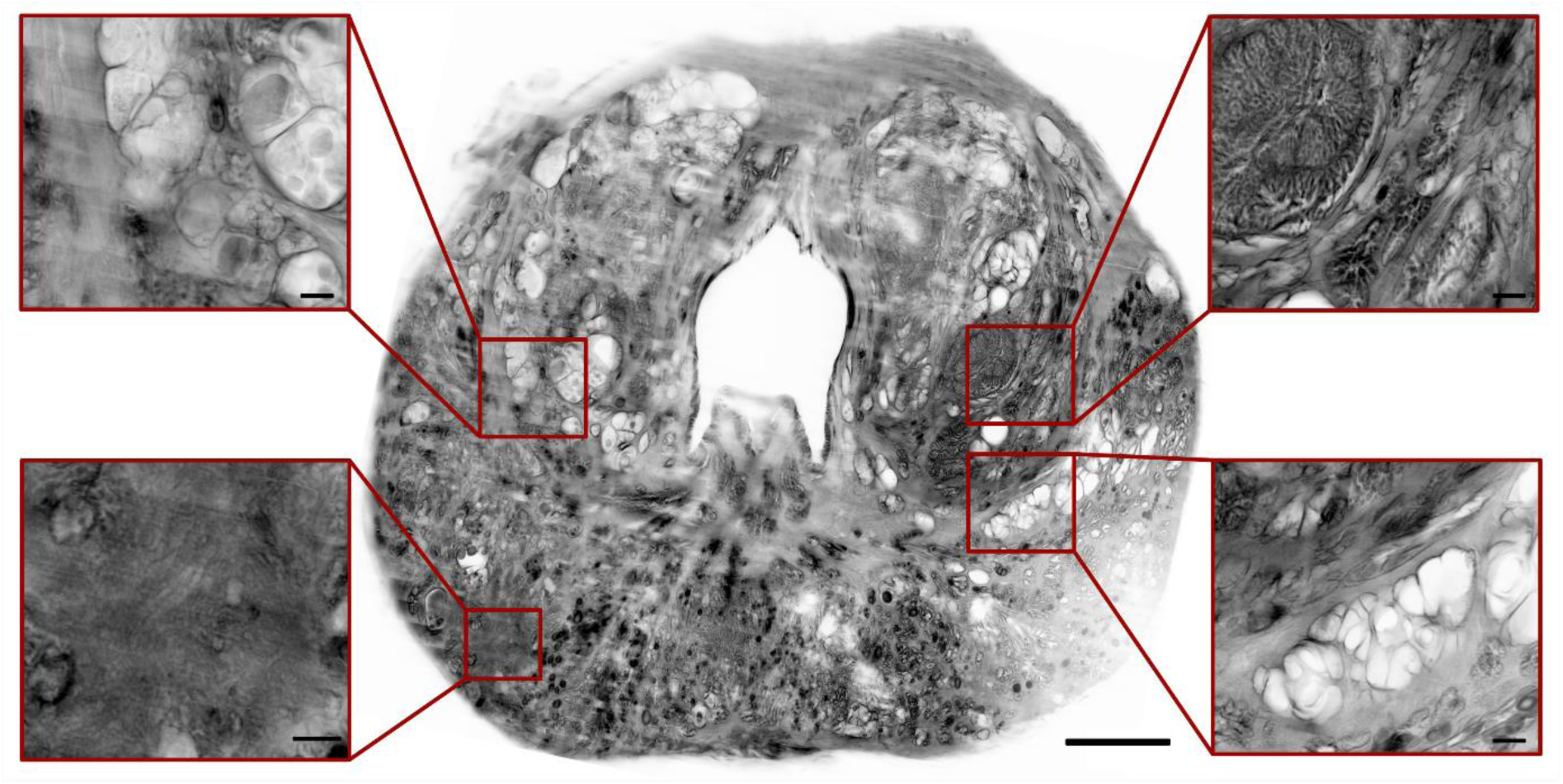
Mosaic 4 acquisition of the entire surface of a prostatectomy section sample. The image shows the stitched surface scan of whole prostate slice at 4 µm isotropic voxel size (middle), with 4 regions highlighted in the magnified inserts. Upper left and lower right: Area of a benign prostatic glandular hyperplasia, consisting of large dilated prostatic glands with intraluminal invaginations. Upper right: Normal prostatic glands of the transition zone consisting of middle large prostatic glands with intervening stroma. Lower left: Tumour region, which appears optically dense in the downsampled 4 µm data set. This is likely due to the many small round oval neoplastic glands with little intervening stroma and without intraluminal projections that make up the tumour, as was confirmed by histopathological evaluation of the corresponding H&E section. Scale bars: Whole sample scan: 5 mm; inserts: 500 µm, respectively.

**Supplementary figure 5:**
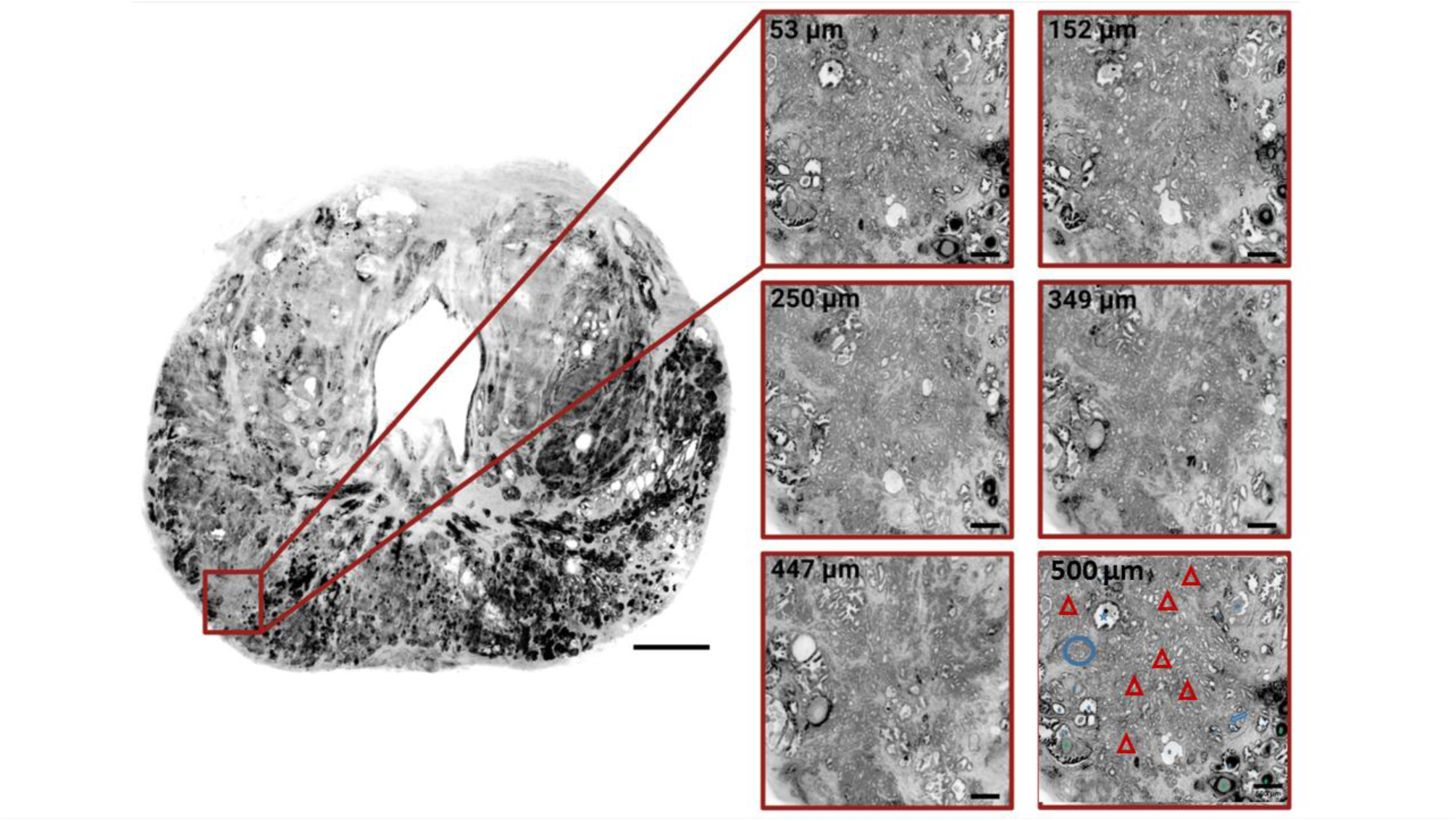
Changes of tumour morphology over depth in 3D. Left: 3D rendering of the entire prostate volume, with the tumour highlighted in red. Right: Tumour morphology at different depths throughout the data volume. The tumour consists of round oval neoplastic glands, compatible with the Gleason pattern 3. In deeper levels (500um) the neoplastic glands are packed very closed to each other, and fusion of some glands might be considered (blue circle), compatible with the higher grade of the prostate cancer grading system, namely Gleason pattern 4. Other area showed a focus of a small glands with almost inconspicuous lumen (blue arrow), morphologically might be compatible with the poorly formed glands of Gleason grade 4. Scale bars: 5 mm (left); 500 µm (right, all panels). Benign prostatic glands can be recognised by irregular gland surface and larger volume (blue stars), as compared with the neoplastic proliferation (red triangles). Corpora amylacea are easy identifiable in the benign glands (green stars).

## References

1. Munck, S., et al., Maximizing content across scales: Moving multimodal microscopy and mesoscopy toward molecular imaging. Curr Opin Chem Biol, 2021. 63: p. 188–199.

2. Markram, H., Reconstruction and Simulation of Neocortical Microcircuitry. Cell, Oktober 2015. 163.

3. Epstein, J.I., et al., The 2014 International Society of Urological Pathology (ISUP) Consensus Conference on Gleason Grading of Prostatic Carcinoma: Definition of Grading Patterns and Proposal for a New Grading System. Am J Surg Pathol, 2016. 40(2): p. 244–52.

4. Nir, G., et al., Automatic grading of prostate cancer in digitized histopathology images: Learning from multiple experts. Med Image Anal, 2018. 50: p. 167–180.

5. Paulk, A.T., I.A. Sesterhenn, and A.P. Burke, Recutting Blocks of Prostate Needle Biopsies: How Much Diagnostic Yield Is Gained? Int J Surg Pathol, 2020. 28(5): p. 490–495.

6. Chung, K., et al., Structural and molecular interrogation of intact biological systems. Nature, 2013. 497(7449): p. 332–7.

7. Liebmann, T., et al., Three-Dimensional Study of Alzheimer’s Disease Hallmarks Using the iDISCO Clearing Method. Cell Rep, 2016. 16(4): p. 1138–1152.

8. Liu, A.K., et al., Bringing CLARITY to the human brain: visualization of Lewy pathology in three dimensions. Neuropathol Appl Neurobiol, 2016. 42(6): p. 573–87.

9. Renier, N., et al., Mapping of Brain Activity by Automated Volume Analysis of Immediate Early Genes. Cell, 2016. 165(7): p. 1789–1802.

10. Renier, N., et al., iDISCO: a simple, rapid method to immunolabel large tissue samples for volume imaging. Cell, 2014. 159(4): p. 896–910.

11. Sabdyusheva Litschauer, I., et al., 3D histopathology of human tumours by fast clearing and ultramicroscopy. Scientific Reports, 2020. 10(1): p. 17619.

12. Susaki, E.A., et al., Versatile whole-organ/body staining and imaging based on electrolyte-gel properties of biological tissues. Nat Commun, 2020. 11(1): p. 1982.

13. Pesce, L., et al., 3D molecular phenotyping of cleared human brain tissues with light-sheet fluorescence microscopy. Commun Biol, 2022. 5(1): p. 447.

14. Zhao, S., et al., Cellular and Molecular Probing of Intact Human Organs. Cell, 2020. 180(4): p. 796–812 e19.

15. Pesce, Fast volumetric mapping of human brain slices. bioarchive (preprint), 2020.

16. Costantini, I., et al., A versatile clearing agent for multi-modal brain imaging. Sci Rep, 2015. 5: p. 9808.

17. Migliori, B., et al., Light sheet theta microscopy for rapid high-resolution imaging of large biological samples. BMC Biol, 2018. 16(1): p. 57.

18. Glaser, A.K., et al., Multi-immersion open-top light-sheet microscope for high-throughput imaging of cleared tissues. Nat Commun, 2019. 10(1): p. 2781.

19. !!! INVALID CITATION !!! [13].

20. Guo, M., et al., Rapid image deconvolution and multiview fusion for optical microscopy. Nat Biotechnol, 2020. 38(11): p. 1337–1346.

21. !!! INVALID CITATION !!! [12].

22. Federa, Human Tissue and Medical Research: Code of Conduct for responsible use. Committee for Guidelines in Research (COREON), 2011.

23. Hildebrand, S., et al., Scalable Labeling for Cytoarchitectonic Characterization of Large Optically Cleared Human Neocortex Samples. Sci Rep, 2019. 9(1): p. 10880.

24. McIlvaine, T.C., A buffer solution for colorimetric comparison. Journal of Biological Chemistry, 1921. 49(1): p. 183–186.

25. Kumar, A., et al., Dual-view plane illumination microscopy for rapid and spatially isotropic imaging. Nature Protocols, 2014. 9(11): p. 2555–2573.

26. Edelstein, A.D., et al., Advanced methods of microscope control using μManager software. Journal of Biological Methods, 2014. 1(2): p. e10.

27. Schindelin, J., et al., Fiji: an open-source platform for biological-image analysis. Nat Methods, 2012. 9(7): p. 676–82.

28. Horl, D., et al., BigStitcher: reconstructing high-resolution image datasets of cleared and expanded samples. Nat Methods, 2019. 16(9): p. 870–874.

29. Li, Y., et al., Incorporating the image formation process into deep learning improves network performance. Nat Methods, 2022. 19(11): p. 1427–1437.

30. Voigt, F.F., et al., The mesoSPIM initiative: open-source light-sheet microscopes for imaging cleared tissue. Nat Methods, 2019. 16(11): p. 1105–1108.

31. Chakraborty, T., et al., Converting lateral scanning into axial focusing to speed up three- dimensional microscopy. Light Sci Appl, 2020. 9: p. 165.

32. Herculano-Houzel, S. and R. Lent, Isotropic fractionator: a simple, rapid method for the quantification of total cell and neuron numbers in the brain. J Neurosci, 2005. 25(10): p. 2518–21.

33. Leuba, G. and L.J. Garey, Comparison of neuronal and glial numerical density in primary and secondary visual cortex of man. Experimental Brain Research, 1989. 77(1): p. 31–38.

34. Pan, C., et al., Shrinkage-mediated imaging of entire organs and organisms using uDISCO. Nat Methods, 2016. 13(10): p. 859–67.

35. Rusch, H., et al., Finding the best clearing approach - Towards 3D wide-scale multimodal imaging of aged human brain tissue. Neuroimage, 2022. 247: p. 118832.

36. Mai, H., et al., Scalable tissue labeling and clearing of intact human organs. Nat Protoc, 2022. 17(10): p. 2188–2215.

37. !!! INVALID CITATION !!! [18, 30].

38. Glaser, A.K., et al., Light-sheet microscopy for slide-free non-destructive pathology of large clinical specimens. Nat Biomed Eng, 2017. 1(7).

39. Glaser, A.K., et al., Multi-immersion open-top light-sheet microscope for high-throughput imaging of cleared tissues. Nature Communications, 2019. 10(1): p. 2781.

40. Chakraborty, T., et al., Light-sheet microscopy of cleared tissues with isotropic, subcellular resolution. Nat Methods, 2019. 16(11): p. 1109–1113.

41. Glaser, A.K., et al., A hybrid open-top light-sheet microscope for multi-scale imaging of cleared tissues. bioRxiv, 2021: p. 2020.05.06.081745.

42. Wu, Y., et al., Spatially isotropic four-dimensional imaging with dual-view plane illumination microscopy. Nat Biotechnol, 2013. 31(11): p. 1032–8.

43. Bouchard, M.B., et al., Swept confocally-aligned planar excitation (SCAPE) microscopy for high speed volumetric imaging of behaving organisms. Nat Photonics, 2015. 9(2): p. 113–119.

44. Gao, L., et al., Lattice light sheet microscopy using tiling lattice light sheets. Opt Express, 2019. 27(2): p. 1497–1506.

